# Soils and sediments host novel archaea with divergent monooxygenases implicated in ammonia oxidation

**DOI:** 10.1101/2021.05.03.442362

**Authors:** Spencer Diamond, Adi Lavy, Alexander Crits-Christoph, Paula B. Matheus Carnevali, Allison Sharrar, Kenneth H. Williams, Jillian F. Banfield

**Author notes:** **Corresponding authors**: Spencer Diamond, Associate Project Scientist, Earth and Planetary Science Department, University of California - Berkeley, Innovative Genomics Institute Building, 2151 Berkeley Way, Berkeley, CA 94720, Jillian F. Banfield, Professor, Earth and Planetary Science Department, University of California - Berkeley, Innovative Genomics Institute Building, 2151 Berkeley Way, Berkeley, CA 94720.

## Abstract

Copper membrane monooxygenases (CuMMOs) play critical roles in the global carbon and nitrogen cycles. Organisms harboring these enzymes perform the first, and rate limiting, step in aerobic oxidation of ammonia, methane, or other simple hydrocarbons. Within archaea, only organisms in the order Nitrososphaerales (Thaumarchaeota) encode CuMMOs, which function exclusively as ammonia monooxygenases. From grassland and hillslope soils and aquifer sediments, we identified 20 genomes from distinct archaeal species encoding divergent CuMMO sequences. These archaea are phylogenetically clustered in a previously unnamed Thermoplasmatota order, herein named the Ca. Angelarcheales. The CuMMO proteins in Ca. Angelarcheales are more similar in structure to those in ammonia-oxidizing archaea than those of bacteria, and they contain all functional residues required for activity. Similarly to the Nitrososphaerales, Ca. Angelarcheales genomes are significantly enriched in blue copper proteins (BCPs) relative to sibling lineages, including plastocyanin-like electron carriers and divergent nitrite reductase-like (nirK) 2-domain cupredoxin proteins co-located with electron transport machinery. Angelarcheales do not have identifiable genes for methanol oxidation or carbon fixation, encode significant capacity for peptide/amino acid uptake and degradation, and share numerous electron transport mechanisms with the Nitrososphaerales. In the studied soils and sediments Ca. Angelarcheales were at least as abundant as ammonia-oxidizing Nitrososphaerales. Thus, we predict that Angelarcheales live a mixotrophic lifestyle based on oxidation of ammonia liberated from peptide and amino acid degradation. This work expands the known diversity of Thermoplasmatota and of CuMMO enzymes in archaea and suggests that these organisms are important and previously unaccounted for contributors to nitrogen cycling.

## INTRODUCTION

Copper membrane monooxygenases (CuMMOs) are a family of phylogenetically diverse and ecologically widespread but highly conserved enzymes that function in the aerobic oxidation of ammonia, methane, and potentially other hydrocarbon substrates^1–3^. Organisms encoding CuMMO are critically important in global carbon and nitrogen cycles^4,5^, as particulate methane monooxygenases (pMMOs) attenuate atmospheric methane release^5^, and ammonia monooxygenases (AMOs) can generate nitrous oxide as a byproduct^6^. While the diversity of CuMMOs has been expanded due to cultivation independent studies^7–9^, CuMMOs have still only been identified in a few monophyletic clades of Bacteria and Archaea^1^. The distribution and functionality of CuMMOs in Archaea is particularly constrained. Archaeal CuMMOs have only been shown to participate in ammonia oxidation, and Nitrososphaerales (Thaumarchaeota) are the only identified group to date that encode AMOs^8^. These ammonia oxidizing archaea (AOA) occur across a wide array of natural environments^10^, and are often far more abundant than ammonia oxidizing bacteria (AOB)^11^, resulting in large contributions to the global nitrogen cycle^3,12^.

The CuMMO is a membrane bound protein complex that consists of 3 core proteins amoA/pmoA, amoB/pmoB, and amoC/pmoC, that are typically encoded in an operon^1,13^. In AOA, a fourth hypothetical protein, amoX, of unknown function is also present^13^. Despite their ecological importance, the difficulties in sustaining in vitro activity of CuMMO complexes as well as difficulties in protein structure resolution has led to inconsistent reports regarding the co-factors and substrate specificity controls for these enzymes. Indeed the substrate specificity of CuMMOs is promiscuous and purified enzymes have been demonstrated to have activity on methane, ammonia, and small hydrocarbons^14,15^. However, in the organismal context, the substrate specificity is highly preferential to a single substrate, and occurs in a metabolic context of enzymes that support the processing of that specific substrate and its byproducts^3,16^.

In both AOA and AOB, ammonia oxidation proceeds initially by the conversion of ammonia to hydroxylamine and subsequent oxidation of hydroxylamine to nitrite^17^. While the AMO mediated oxidation of ammonia to hydroxylamine has been established in AOA^18^, the lack of an identifiable hydroxylamine dehydrogenase (Hao) in Archaea has led to the proposal of at least two hypothetical routes from hydroxylamine to nitrite^17,19^. Evidence is mounting that Hao activity in AOA instead relies on a suite of blue copper proteins (BCPs), which are numerous in AOA lineages^16,17,19,20^. Plastocyanin-like electron transport proteins, divergent Cu-containing nitrite reductases (nirK), and other 1 and 2 domain cupredoxin-like proteins have been shown to have correlated transcriptional activity with AMO proteins^21^, and are implicated as substituting for the missing Hao activity.

Identification of novel CuMMO complexes in microbes, and surveys of CuMMOs in natural environments, has primarily relied on primer based nucleic acid amplification of the amoA/pmoA subunit^22^. However, the requirement of sufficient sequence similarity between environmental sequences and known bacterial and archaeal amoA/pmoA subunits precludes the identification of completely novel sequence variants. Further, primer-based studies do not provide information about genome context, which can place CuMMOs in a metabolic context that supports a specific activity. Here, we applied sensitive homology-based methods to *de novo* reconstructed metagenome assembled genomes (MAGs) obtained from soils, aquifer sediment, and deep ocean environments to identify novel CuMMO variants and describe their genomic, phylogenetic and ecological contexts. We propose that a new group of soil- and aquifer-associated archaea contribute to the ecosystem essential and possibly keystone functionality of ammonia oxidation that has previously only been linked to a phylogenetically narrow range of bacteria and archaea.

## RESULTS

### Divergent CuMMOs identified in MAGs recovered from soil and sediment ecosystems

In previous work we identified putative divergent amoA/pmoA homologues in 7 Thermoplasmatota genomes recovered from a Mediterranean grassland soil^23^. This was intriguing, given that amo/pmo homologues had not been previously observed in archaea outside of the Nitrososphaerales (Thaumarchaeota). Here we searched for additional genomes encoding related (divergent) amo/pmos using a series of readily available, and custom built, hidden markov models (HMMs) across all archaeal genomes in the Genome Taxonomy Database (GTDB), and in all archaeal MAGs in our unpublished datasets from ongoing studies (**Supplementary Fig. 1**). We found additional amoA/pmoA genes in genomes recovered from soils at the South Meadow and Rivendell sites of the Angelo Coast Range Reserve (CA)^23,24^, the nearby Sagehorn site^24^, a hillslope of the East River watershed (CO)^25^, and in sediments from the Rifle aquifer (CO)^26^ and the deep ocean^27^. In total we identified 201 archaeal MAGs taxonomically placed using phylogenetically informative single copy marker genes outside of Nitrososphaerales containing divergent amo/pmo proteins (**Supplementary Table 1 and Supplementary Data 1)**. Genome de-replication with additional references at 95% average nucleotide identity (ANI) resulted in 34 species-level genome clusters, 20 of which encoded an amo/pmo homologue (**Supplementary Table 2**). Of these genomes, 11 are species not previously available in public databases. In all cases where assembled sequences were of sufficient length, the amoA/pmoA, B, and C protein coding genes were found co-located with each other and with a hypothetical protein considered to be amoX in the order C-A-X-B (**Fig. 1a, Supplementary Table 2, and Supplementary Fig. 2**). The mean sequence identity of the recovered amoA/pmoA, B, and C proteins to known bacterial sequences were 16.7, 8.0, and 14.2 % and 13.8, 9.5, and 20.8 % to known archaeal sequences. As might be expected considering the large sequence divergence between the recovered sequences and known amo/pmo proteins, we found that no pair of typical primers used for bacterial and archaeal amoA/pmoA environmental surveys^28^ matched any novel amoA/pmoA gene with less than 7 mismatched bases (**Supplementary Table 3**). This suggests that these sequences would have been missed by previous primer-based pmoA gene surveys.

**Figure 1.**
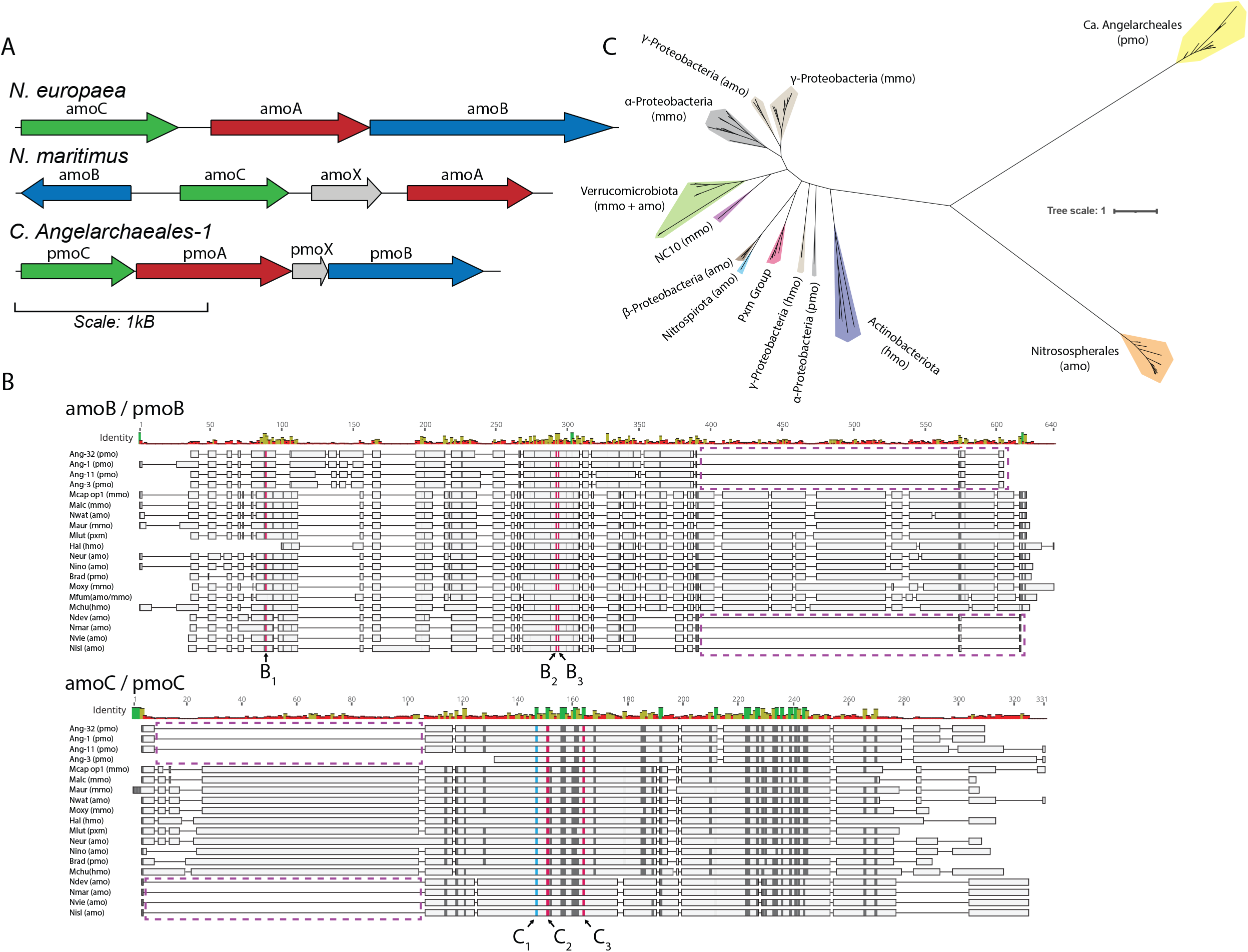
Comparison of divergent CuMMO proteins to known sequences. **(A)** Comparison of pmoABC loci and gene arrangement between Nitrosomonas europaea (ammonia oxidizing bacteria), Nitrosopumilus maritimus (ammonia oxidizing archaea), and Ca. Angelarcheales-1. **(B)** Representative sequences from a multiple sequence alignment of amoB/pmoB proteins (top; n = 114 sequences) and amoC/pmoC proteins (bottom; n = 110 sequences). Sequences are named by species abbreviation followed by putative function in parentheses: pmo = particulate monooxygenase (no biochemical evidence), mmo = methane monooxygenase, amo = ammonia monooxygenase, hmo = hydrocarbon monooxygenase. Species abbreviations are described in Supplementary Table 3. Lettering below alignments indicates residues forming part of B or C-sites in these proteins. Histidine residues and aspartic acid residues within these sites are colored red and blue respectively. Residues colored black are identical across >= 65% of sequences in full alignment. Purple boxes indicate truncations shared by CuMMOs identified in this study and known archaeal sequences. All amoABC/pmoABC proteins and alignments are available in Supplementary Data 1. **(C)** Maximum likelihood phylogenetic tree constructed from a concatenated alignment of amoABC/pmoABC proteins (n = 112 sequences; >= 2 sequences per genome). Colored clades are drawn based on a combination of shared function and taxonomy and colored by the taxonomy of genomes encoding the sequences within the clade. Clades are labeled by their shared taxonomy and known/predicted CuMMO function in parentheses (see above). Pxm group is an exception and represents a group of duplicated proteins present in many gammaproteobacterial methanotrophs. Scale bar represents average changes per amino acid position.

### Novel CuMMO subunit sequences contain expected catalytic and metal binding residues

Alignments were constructed for each predicted amoA/pmoA, B, and C protein in combination with reference sequences that cover the known diversity of these protein subunits^1^ (**Fig. 1b and Supplementary Data 1**). In the new sequences, all of the expected catalytic and metal binding residues^29^ were conserved (**Fig. 1b**). In amoB/pmoB, all three histidine residues for the mono-copper B-site required for enzyme activity were conserved. The C-site in amoC/pmoC, which contains an aspartic acid and two histidine residues important for enzymatic activity^13^, is also completely conserved. Although the A-sites in amoB/pmoB and the D-sites in amoC/pmoC and amoA/pmoA were not observed in the new sequences, these sites are not required for catalytic function and are only conserved within bacterial lineages^1,8^. We also note that the new amoB/pmoB sequences share a C-terminal truncation to previously identified Nitrososphaerales pmoB’s^30,31^, as well as share an N-terminal truncation in the amoC/pmoC protein.

### Novel CuMMOs form a new group in the CuMMO superfamily

To infer the evolutionary relationship of our newly identified CuMMO sequences to known CuMMO family members we used a concatenated pmoABC sequence alignment to produce a phylogenetic reconstruction covering known family members **(Fig. 1c)**. Individual protein phylogenies were also constructed for each pmo subunit (**Supplementary Fig. 3a-c**). The different subunit reconstructions largely agree in overall topology, and all reconstructions support our newly identified sequences as a highly divergent third major lineage of CuMMOs. Also, similar to previous phylogenetic reconstructions of CuMMO sequences, our sequences form clusters that mirror the phylogenetic relatedness of encoding genomes^26^.

### Archaea with divergent CuMMOs form a novel clade within the Thermoplasmatota

An initial phylogenetic classification of our archaeal genomes placed them as an order level lineage, RBG-16-18-12, within the Thermoplasmatota phylum (**Supplementary Tables 1-3**). A concatenated alignment of 76 archaeal specific marker proteins^32^ resolved them as a well supported order-level monophyletic lineage basal to the *Methanomassilococalles* within *Thermoplasmatota* (**Fig. 2a, Supplementary Fig. 4, and Supplementary Table 4**). We reported the first genomes from this clade (RBG-16-68-12 in GTDB) from the RBG dataset from the Rifle, CO aquifer sediments^23^. Given that this clade is now represented by 32 species-level genomes (with 11 additional species added in this study), and that 10 genomes satisfy the completeness and contamination requirements to be considered high-quality drafts^1,8^, we propose that they define a new candidate order, hereafter referred to as the Ca. Angelarcheales. We propose Ca. Angelarcheales-1 (GCA_005878985.1^16^) from the Angelo Coastal Reserve, CA, as the type genome for this clade. Of the Ca. Angelarcheales genome set, 20 contain identifiable pmo/amo gene clusters. The fact that CuMMO sequences are encoded within genomes of a single monophyletic subclade of the Thermoplasmatota is similar to previous observed patterns of CuMMO distribution, as taxa that encode CuMMO systems are often constrained to monophyletic groups scattered across the tree of life^3,17,19^.

**Figure 2.**
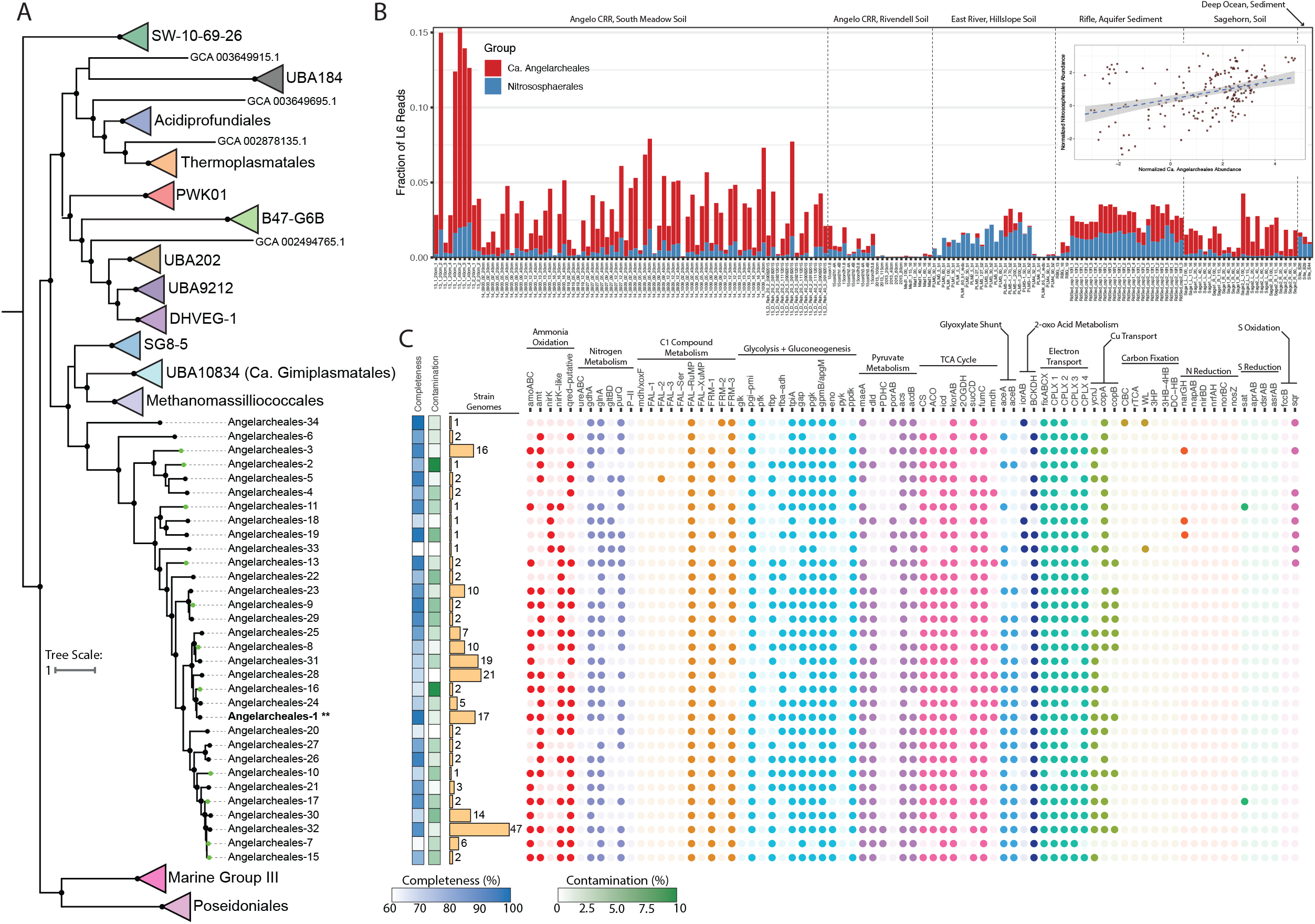
Taxonomy, metabolism, and distribution of Ca. Angelarcheales. **(A)** Maximum likelihood phylogenetic tree constructed for the archaeal phylum Thermoplasmatota using a concatenated alignment of 76 archaeal specific marker genes. The tree includes 32 Ca. Angelarcheales genomes and 295 reference genomes. The type strain, Ca. Angelarcheales-1, is marked with two asterisks. Tree was rooted using A. fulgidus (GCF_000008665.1) as an outgroup. Clades were collapsed at the order level if they contained more than one genome and arbitrarily colored. Black dots at nodes indicate >= 95% bootstrap support (ufboot; n = 1000). For the full, un-collapsed, tree see Supplementary Fig. 4. **(B)** Relative abundance information, based on rpL6 read counts, for Ca. Angelarcheales and Nitrososphaerales across 185 shotgun metagenome samples from 6 sites. The x-axis indicates the sample name, and the y-axis indicates the fraction of reads out of the total reads in a given sample that mapped to rpL6 sequences taxonomically associated with each group. Samples are separated by the general sampling location, indicated at the top of the plot. Inset, normalized rpL6 based relative abundance of Ca. Angelarcheales (x-axis) vs Nitrososphaerales (y-axis) for all 185 shotgun metagenome samples. A best fit line is plotted using linear regression, shaded area indicates standard error of the regression. Rho of association is positive and significant (rho = 0.366, FDR < 0.001). **(C)** Genome quality, number of strain-level genomes (genomes with ANI >= 95%), and predicted metabolism for the 32 Ca. Angelarcheales genomes in Fig. 2A. Filled dots indicate the presence of a gene or gene set that executes a specific metabolic function or reaction. Dots are colored based on shared pathways or metabolic functionality as described above the figure, and colors are chosen arbitrarily. For complete explanation of metabolic functions and search criteria see Supplementary Table 8.

### Taxonomic marker profiling indicates widespread distribution of Ca. Angelarcheales

Using ribosomal protein L6 (rpL6) as a taxonomic marker, we determined an average prokaryotic community relative abundance of 1.73 ± 2.25 % for Ca. Angelarcheales in 185 samples from six environments **(Fig. 2b and Supplementary Tables 5-6)**. In comparison, the average relative abundance of Nitrososphaerales in the same dataset was 0.65 ± 0.61 %. We observed significant and strong positive associations between the abundances of both Ca. Angelarcheales and Nitrososphaerales with each other (**Fig. 2b;** rho = 0.366, FDR < 0.001), and with the bacterial order Nitrospirales (rho_Ang_ = 0.400, FDR < 0.001; rho_Nitroso_ = 0.613; FDR < 0.001), a group of nitrite reducing bacteria (**Supplementary Fig. 5 and Supplementary Table 7**).

### Ca. Angelarcheales share similar metabolic features with ammonia oxidizing Nitrososphaerales

We conducted a general metabolic analysis of Ca. Angelarcheales to place the CuMMO into metabolic context (**Fig. 2c, Supplementary Fig. 6a-d, Supplementary Table 8, and Supplementary Data 2**). Ca. Angelarcheales genomes contained many of the electron transfer and ammonia assimilation components known to be conserved in characterized ammonia oxidizing archaea including: an NADH:ubiquinone oxidoreductase complex with an additional copy of the M protein component (CPLX 1), a four-subunit putative succinate dehydrogenase complex (CPLX II), a complete cytochrome b containing complex III with a plastoquinone-like electron transfer apparatus (CPLX 3), up to two distinct oxygen reducing terminal oxidases (CPLX 4), an ammonia transporter (amt), a glutamine synthase (glnA), and glutamate dehydrogenase (gdhA). We found little evidence that Ca. Angelarcheales can use inorganic nitrogen or sulfur containing compounds as alternative electron acceptors, thus it is likely that these organisms are obligate aerobes. We also did not identify carbon fixation pathways within any CuMMO encoding Ca. Angelarcheales.

Some genomes encode a credible nitrite reductase (nirK) and 2-domain cupredoxins with homology to nirK (nirK-like). However, nirK is not essential for ammonia oxidation in Nitrososphaerales^33^. Also identified were plastocyanin-like proteins, which are common in Nitrososphaerales (**Fig. 3)**, and three distinct copper transport systems (**Fig. 2c**). The parallels between the redox enzyme inventories of Ca. Angelarcheales and Nitrososphaerales support the role of the CuMMO in ammonia rather than methane oxidation. Further supporting this deduction, we did not detect any genes with homology to methanol dehydrogenases (mdh/xoxF) across the entire Ca. Angelarcheales clade, which would be expected if methane to methanol oxidation was occurring. Mechanisms to assimilate formaldehyde and formate (breakdown products of methanol) were present, however these compounds can originate from other processes, such as sarcosine and betaine degradation pathways, which are present in Ca. Angelarcheales.

**Figure 3.**
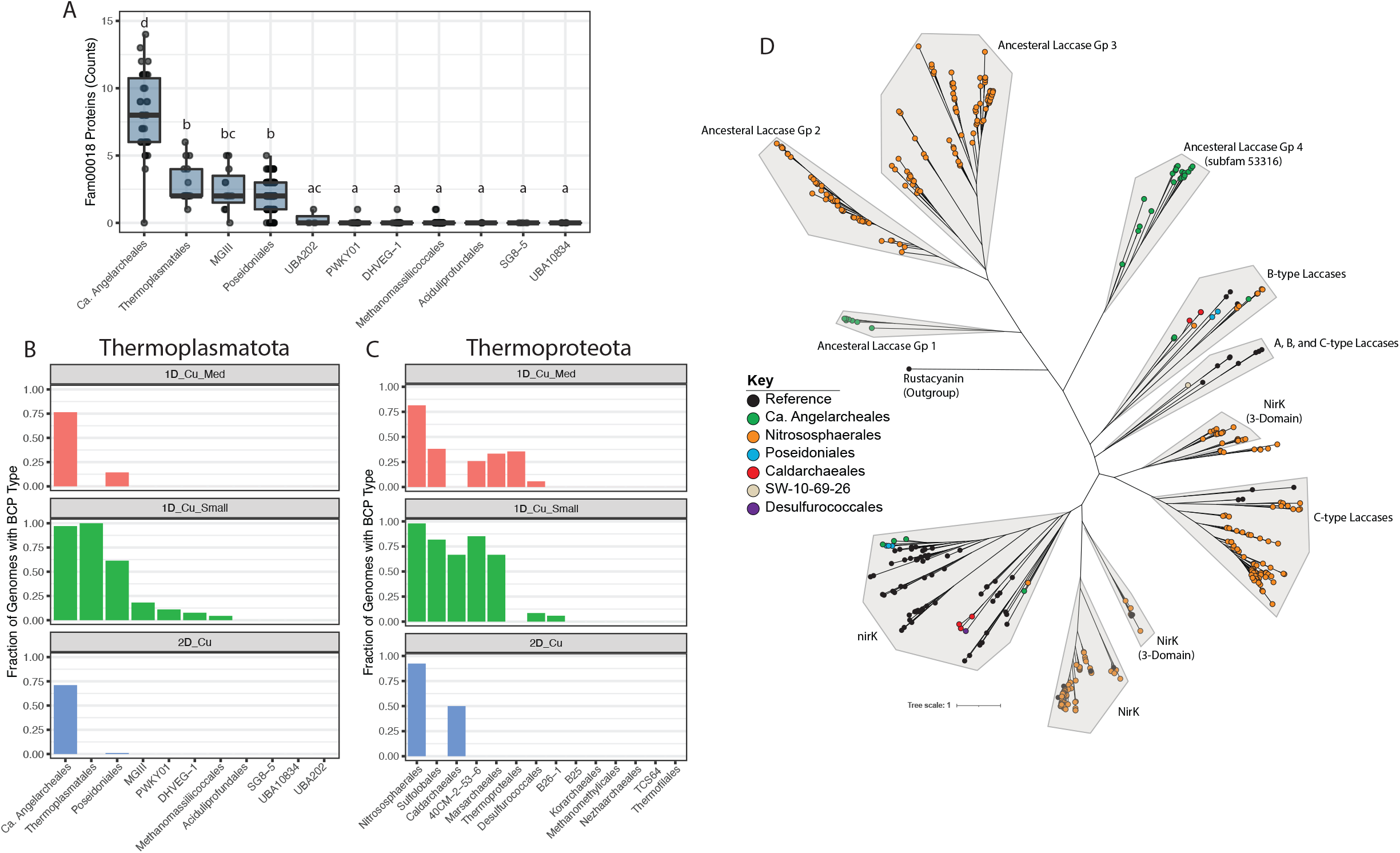
Analysis of BCPs across CuMMO encoding archaeal phyla. **(A)** Counts of fam00018 proteins in genomes from each order level lineage of the Thermoplasmatota containing ≥ 3 genomes. Each dot represents one genome. Boxes indicate the first and third quartile of counts, lines in boxes indicate median values, and whiskers indicate 1.5 x IQR in either direction. Letters above boxes indicate statistically significant differences between groups. Groups sharing no letters have statistically significant differences (FDR ≤ 0.05; pairwise Wilcoxon test). **(B)** The fraction of genomes within each order level lineage of the Thermoplasmatota containing ≥ 3 genomes carrying the BCP subtype noted in the plot title. **(C)** The fraction of genomes within each order level lineage of the Thermoproteota containing ≥ 3 genomes carrying the BCP subtype noted in the plot title. **(D)** Maximum likelihood phylogenetic tree containing 349 2-domain and 3-domain BCPs from our analysis and 90 reference nirK and 2-domain laccase proteins. Clades were manually defined, shaded in grey, and named based on their constituent reference sequences or based on their sequence architecture relative to expected ancestral 2-domain BCPs34. Node colors indicate if a sequence was a reference sequence (black) or the order level taxonomy of the encoding genome. Rustacyanin (ACK80662.1) is provided as an outgroup.

Ca. Angelarcheales contain numerous transporters for branched chain amino acids, polar and non-polar amino acids, oligopeptides, and many proteases (**Supplementary Fig. 6b-d**). The number of encoded amino acid and peptide transport systems in Ca. Angelarcheales is on average the largest across the phylum Thermoplasmatota (**Supplementary Fig. 6d**). The presence of a branched chain keto acid dehydrogenase complex (BCKDH) enables the degradation of branched chain amino acids to acetyl and propionyl-CoA, and a glyoxylate shunt (aceA and aceB) enables the carbons of acetyl-CoA to be used for biosynthesis. A number of enzymes indicate the capacity for acetate degradation to acetyl-CoA (acetate-CoA ligase (acdB)) and lactate degradation to pyruvate (D-lactate dehydrogenase (dld)). These archaea do not have a complete glycolytic pathway (missing core enzymes including glucokinase (glk), phosphofructokinase (pfk), and pyruvate kinase (pfk)), but have gluconeogenesis pathways, thus enabling the biosynthesis of glucose from acetyl-CoA and pyruvate.

### Blue copper proteins (BCPs) are enriched in CuMMO containing archaea

As BCPs have been suggested to serve critical functions in archaeal ammonia oxidation^16^, we compared the BCP inventories in genomes of the phyla Thermoplasmatota and Thermoproteota, which include Ca. Angelarcheales and Nitrososphaerales, respectively. The dataset included 34 representative Ca. Angelarcheales genomes and 610 reference genomes (**Supplementary Table 4**). Due to their high primary sequence diversity, the identification and comparison of BCPs across organisms is difficult using standard annotation methods. Thus we clustered 1,103,913 proteins using a previously validated two-step protein clustering approach^33^. This generated 76,216 protein subfamily clusters (subfams), which are groups of proteins sharing global homology, and 19,828 protein family clusters (fams), which are groups of protein subfamilies where remote local homology could be confidently detected. We identified 1,927 proteins with BCP-associated (cupredoxin-like) PFAM domains across 30 protein fams (**Supplementary Fig. 7a**). Notably, a single protein family (fam00018) contained 1,738 (90.2%) of these proteins, and the remaining proteins either made up very small fractions of other fams or were part of fams with very few proteins (**Supplementary Fig. 7b**). Analysis of the domain architectures of proteins within fam00018 indicate that this protein family primarily contains BCPs with between 1-3 cupredoxin-like domains. Included in fam00018 are small globular plastocyanin-like proteins, nirK-like proteins, 2-domain laccase-like proteins, and the Cu binding cytochrome c oxidase subunit 2 (coxB/COX2) (**Supplementary Fig. 7c**). Fam00018 also contained 671 proteins with no identifiable domain annotations, which was expected given the high sequence diversity of BCPs. However, many proteins with no annotations were clustered into fam00018 subfamilies containing proteins with identifiable BCP domains, allowing the recruitment of these proteins into our analyses. We used the proteins of fam00018 as a broad homology group to quantify and ultimately sub-classify BCP types across genomes (**Supplementary Table 9**). Compared to all other order-level lineages in the Thermoplasmatota, Ca. Angelarcheales genomes are significantly enriched in fam00018 proteins (FDR < 0.05 ; pairwise Wilcoxon test), encoding on average 8.1 per genome **(Fig. 3a)**. This pattern of fam00018 protein enrichment is similarly observed for the ammonia-oxidizing Nitrososphaerales order relative to sibling orders within the Thermoproteota (FDR < 0.05 ; pairwise Wilcoxon test) (**Supplementary Fig. 7d)**.

### Subclassification of fam00018 identifies specific BCP architectures associated with lineages carrying CuMMOs

To more comprehensively understand the subtypes of BCPs that are present across the archaeal orders within Thermoplasmatota and Thermoproteota, we subdivided fam00018 into six manually annotated groups that covered 85.3% of all fam00018 proteins (**Fig. 3b, 3c, Supplementary Figs. 8a-b, and Supplementary Table 9**). We observed that small plastocyanin-like 1-domain BCPs (< 250 aa), while present in many lineages, were extremely prevalent in the genomes of Ca. Angelarcheales and Nitrososphaerales, supporting their important role in facilitating electron transport in these groups (**Figs. 3b and 3c**). Alternatively, medium length 1-domain BCPs (250-400 aa) and 2-domain BCPs were encoded by most genomes of Nitrososphaerales and Ca. Angelarcheales, found in few lineages outside them, and if found were not widely present in the genomes of those other lineages (**Figs. 3b and 3c**). This is consistent with these proteins performing functions that are specific to both Ca. Angelarcheales and Nitrososphaerales.

We speculated that two-domain cupredoxins (2-domain BCPs) generally, and not nirK specifically, may be important in the ammonia oxidation pathway in archaea. These proteins can be differentiated based on phylogenetic relationships, the types of copper centers they contain, and the arrangement of these centers^34,35^. A phylogenetic tree for 2-domain BCPs, known nirK sequences (which include 3-domain BCPs), and 2-domain laccase sequences **(Fig. 3d)** resolves 11 discrete clades. The Nitrososphaerales nirK clades are distinct from classic nirK sequences, as has been observed previously^36^. The four, high confidence, nirK sequences identified in Ca. Angelarcheales fall into the classic nirK clade. Four clades are composed of sequences that contain two Type I copper centers but appear to lack Type II or III centers. Such proteins lack functional predictions, and are referred to as ancestral forms of 2-domain BCPs^34^. The 2-domain BCPs of ancestral group 4 (subfam53316) and ancestral group 1 (subfam54500) are exclusively found in Ca. Angelarcheales. We also note that while 2-domain BCPs were found in archaeal orders outside of Ca. Angelarcheales and Nitrososphaerales, these sequences fall into clades of known laccases.

### BCPs in Ca. Angelarcheales are co-localized with energy generation machinery

We examined the genomic context, in Angelarcheales, of three gene clusters known to be important for electron transfer and energy generation in ammonia oxidizing archaea: the amoCAXB cluster, the coxAB oxygen utilizing terminal oxidase cluster, and the complex III like cytochrome b gene cluster (**Fig. 4a and Supplementary Figs. 2, 9, and 10)**. Four of the amoCAXB encoding contigs from 20 genomes encode a medium length 1-domain BCP from subfam17112 ∼4 genes upstream of the amoCAXB gene cluster. We note that only 5 of 20 contigs have sufficient length upstream of the amoCAXB locus to allow identification of this BCP. The proteins of subfam17112 are all predicted to contain 5 transmembrane helices in their N-terminal region with a cupredoxin-like domain occupying the outer membrane facing C-terminal region (**Supplementary Fig. 11**). The 8 proteins of subfam17112 only occur in Ca. Angelarcheales genomes that also encode an amo. Medium length 1-domain BCPs also occur at a high frequency in Nitrososphaerales (**Fig. 3c**) and have been proposed to be associated with hao activity^16^. Thus, subfam17112 may be implicated in hao activity, given its proximity to the amoCAXB locus and the proposal that an archaeal hao would be a divergent membrane bound cupredoxin-like protein^16,37^.

**Figure 4.**
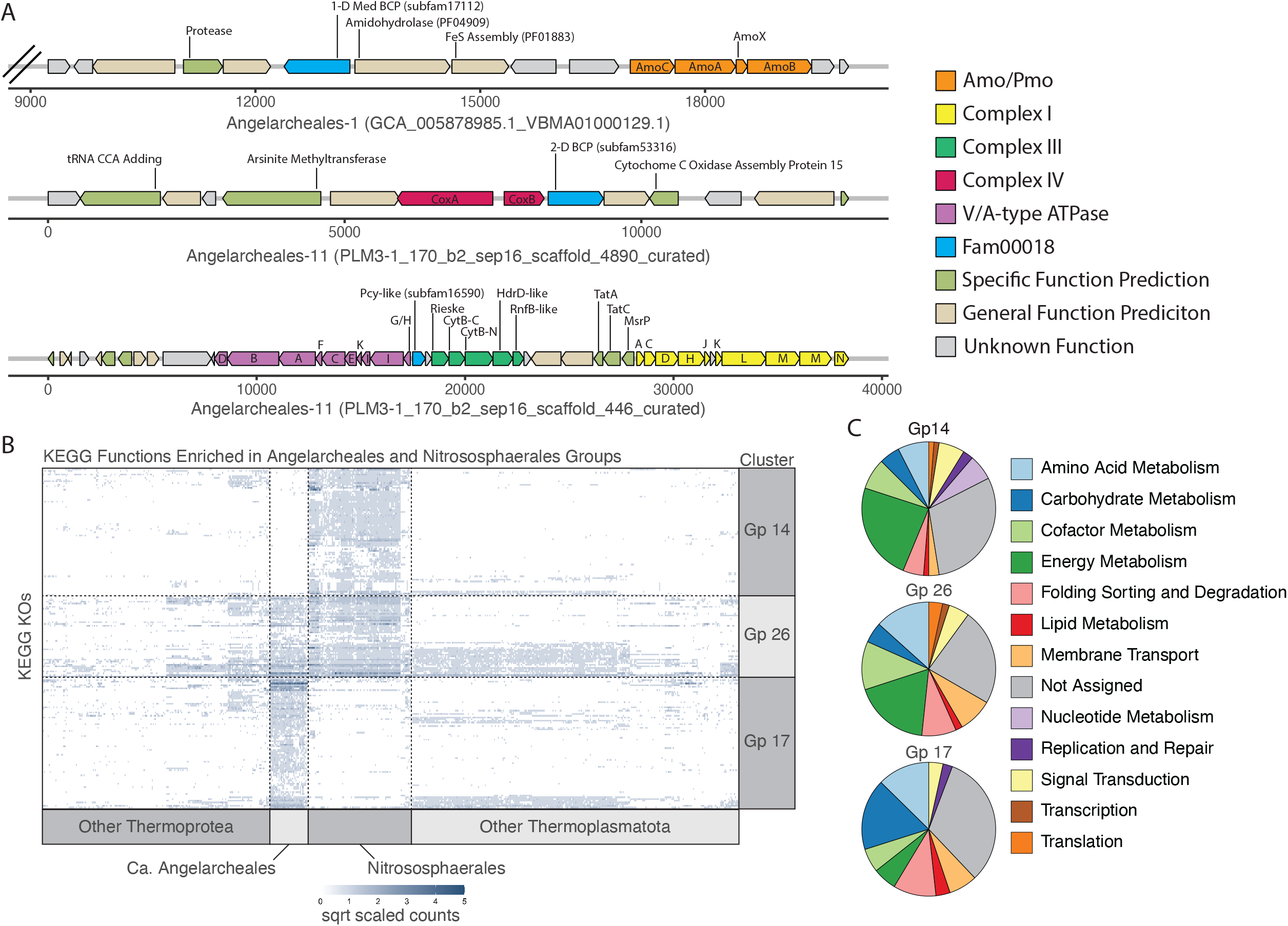
Genomic context and metabolic function enrichment analysis for Angelarcheales. **(A)** Example operons showing the gene context surrounding the amoCAXB gene cluster (top), the coxAB oxygen utilizing terminal oxidase gene cluster (middle), and the complex III like cytochrome b gene cluster (bottom). Subfam membership is noted in labels for fam00018 proteins (blue). Scale is in base pairs and double hash on amoCAXB containing contig indicates truncation after 10 genes to the left of the cluster for readability. **(B)** Heatmap showing the number of hits to 201 KEGG KOs found to be significantly enriched in Angelarcheales (Gp 17), the Nitrososphaerales (Gp 14), or showing shared enrichment by both orders relative to all others (Gp 26). Each column represents the hits across one genome, and each row represents the hits for a single KO. Intensity of each spot in the heatmap is based on square root scaled counts of hits to a KO in each genome for ease of readability. Dotted lines are added to segregate clusters for ease of viewing. **(C)** Breakdowns of functional categories associated with KEGG KOs in each enrichment group. Also see Supplementary Table 10.

We reconstructed coxAB encoding contigs from 29 genomes. In 19 of these genomes, we could identify a 2-domain BCP directly following the coxB gene, which in 14 of 19 cases was the 2-domain BCP from the nirK-like group subfam53316 (**Fig. 3d)**. Again, in at least 7 cases it was not possible to search for a BCP as contigs were of insufficient length.

A cytochrome b containing complex III-like locus could be identified and reconstructed in 30 genomes. It was commonly co-located with gene clusters encoding other components of the electron transport chain (the V/A-type ATPase and NADPH:Quinone oxidoreductase - complex I). Electron transfer from complex III to downstream electron transport machinery is posited to involve a plastocyanin-like 1 domain BCP, not a soluble cytochrome c, similar to ammonia oxidizing Nitrososphaerales^16,37^. In Ca. Angelarcheales, we found a small 1 domain plastocyanin-like BCP gene upstream of a Rieske iron-sulfur protein in 23 of 30 reconstructed complex III loci (and no cytochrome c). This indicates that the electron transport chain in Ca. Angelarcheales is similar to that found in Nitrososphaerales.

### Metabolic functions enriched in Ca. Angelarcheales and Nitrososphaerales

Using indicator analysis, we identified KEGG orthology groups (KOs) that were significantly enriched in the Ca. Angelarcheales (Gp 17), the Nitrososphaerales (Gp 14), and the KOs shared by both orders relative to all other orders of Thermoplasmatota and Thermoproteota (Gp 26). Of the 78 KOs that were significantly enriched in Ca. Angelarcheales (**Fig. 4b and Supplementary Table 10**), the largest functional groups corresponded to carbohydrate metabolism (17.2%), amino acid metabolism (12.6%), and protein folding, sorting, and degradation (10.3%) (**Fig. 4c**). KOs enriched in Ca. Angelarcheales support the use of peptides and amino acids as a carbon and nitrogen source. These included isocitrate lyase of the glyoxylate shunt (K01637), proteins for detoxification of the threonine catabolite methylglyoxal (K10759, K18930, and K23257), 4 proteases (K01392, K06013, K07263, and K09640), components of the archaeal proteosome (K13527 and K13571) enzymes for betaine (K00130, K00544, and K00479), proline (K00318), and cysteine (K01760) catabolism, the E1 component of the branched chain keto acid dehydrogenase complex (K00166), and a transport system for polar amino acids (K02028 and K02029).

The 75 KOs significantly enriched in Nitrososphaerales genomes were largely associated with energy metabolism (23.7%) (**Fig. 4c**). This included functions critical for the hydroxypropionate/hydroxybutyrate carbon fixation pathway known to operate in these organisms (K18593, K18594, K18603, and K18604). The absence of these genes in the Ca. Angelarcheales, thus their inability to fix CO_2_, differentiates them from the Nitrososphaerales. Other functions enriched in Nitrososphaerales are involved in electron transfer including plastocyanins (K02638), ferredoxins (K05524), and rieske iron-sulfur proteins (K15878). We also identified enriched capacity for urea utilization (K01429, K01430, K03187, K03188, K03190) and urea transport (K20989), which agrees with the fact that many Nitrososphaerales are thought to use urea as a nitrogen source for ammonia oxidation^38^.

The AMO subunits A and C (K10944 and K10946) were identified among the 48 functions that were significantly enriched in both Ca. Angelarcheales and Nitrososphaerales compared to the other groups. Many shared functions were also associated with energy metabolism and are known to be associated with ammonia oxidizing archaea, including nitrite reductase (K00368), the oxygen utilizing terminal oxidase subunit I (K02274), the ammonium transporter (K03320), a duplicated NADPH:Quinone oxidoreductase subunit M (K00342), and a split cytochrome b-561 like protein (K15879), as well as 3 iron-sulfur complex assembly proteins (K09014, K09015, and K13628), a cytochrome c oxidase complex assembly protein (K02259), a high affinity iron transporter (K07243), and a copper transporter (K14166).

### Metabolic reconstruction supports an amino acid based metabolism

We undertook a complete metabolic reconstruction for the type strain genome (Angelarchaeales-1) to evaluate the feasibility of a life strategy where amino acid metabolism is coupled to ammonia oxidation. We focused on pathways for the import and catabolism of amino acids and routes by which their products feed into central carbon metabolism, ammonia oxidation, and are interconnected with electron transport and energy generation (**Fig. 5 and Supplementary Tables 11-12)**.

**Figure 5.**
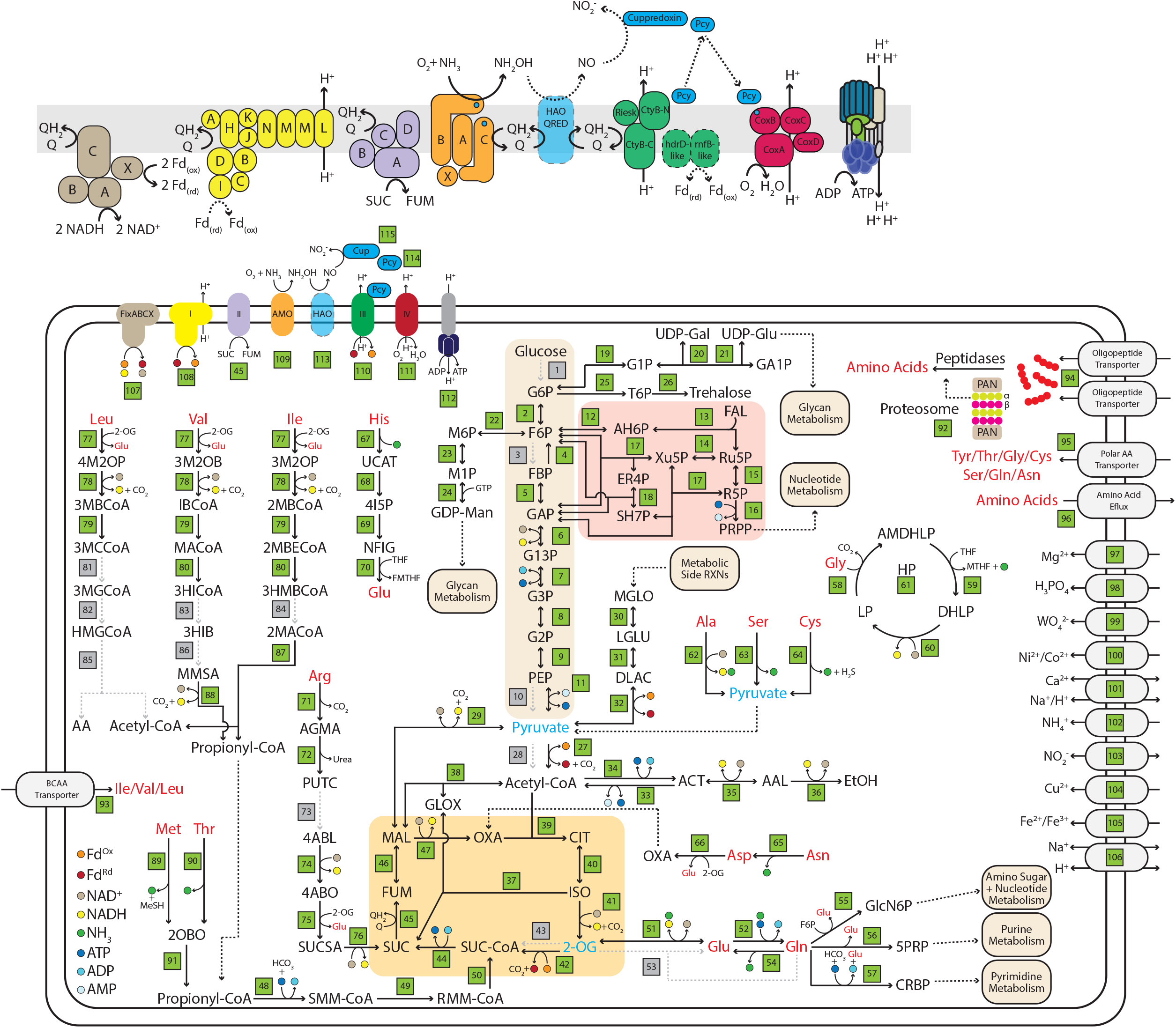
Metabolic reconstruction of Angelarcheales-1 genome. For full reaction, gene list, and list of compound abbreviations see Supplementary Tables 12 and 13. Green boxes indicate a reaction (and its reference number in Supplementary Table 12) that could be linked to a gene with the predicted metabolic function. Black arrows with solid lines indicate a reaction that could be identified, associated smaller arrows with colored dots indicate consumed, and generated reaction substrates and products are indicated in the key at the bottom left. Black arrows with dotted lines indicate flow of metabolites to other pathways or reactions. Grey arrows with dotted lines and grey boxes indicate reactions that were searched for and could not be identified. Amino acids are in red text to highlight their locations throughout the figure. Metabolites in blue text indicate hubs for carbon derived from amino acid catabolism. For ease of viewing the reactions of glycolysis, the pentose phosphate pathway, and the TCA cycle have been highlighted with beige, red, and orange backgrounds. The upper panel is a blow up of the electron transport reactions showing predicted organizations of subunits in each complex. Reference numbers for each subunit can be found in the larger figure panel, and colors of subunits are the same as those used in that figure panel. Transparent HAO-QRED indicates a putative/proposed functionality. Black arrows with dotted lines indicate putative reactions. Protein subunits with solid color but dotted borders indicate a protein was found but functionality is unclear.

As is typical of the Angelarcheales, the Angelarchaeales-1 genome lacks key oxidative enzymes of glycolysis and oxidative enzymes of the pentose phosphate pathway. They can interconvert glucose/fructose to mannose and galactose derivatives but appear unable to import or phosphorylate these sugars. The encoded fructose bisphosphatase would allow gluconeogenesis. They have no detectable pyruvate kinase and instead encode a pyruvate orthophosphate dikinase, which is known to be allosterically regulated and reversible in archaea^39,40^. The capacity for production of the compatible solute trehalose is notable, as Angelarchaeales-1 derives from an environment that regularly undergoes large cyclic changes in water content^23^.

Angelarchaeales-1 encodes a full complement of genes for the conversion of pyruvate into Acetyl-CoA, the TCA cycle, and a glyoxylate shunt. However, it lacks genes for both the pyruvate dehydrogenase complex as well as the 2-oxoglutarate dehydrogenase complex. These reactions are likely enabled by pyruvate/2-oxoglutarate ferredoxin oxidoreductase systems, which provide reduced ferredoxin. A glyoxylate shunt allows for catabolic reactions that terminate in 2-carbon compounds (e.g. acetyl-CoA produced by amino acid and acetate catabolism) to be utilized for biosynthetic purposes, as it bypasses the two decarboxylation steps of the TCA cycle.

We identified reasonably confident catabolic routes for 15 amino acids, including a complete glycine cleavage system, as well as a route for the end product (propionyl-CoA) of at least 4 amino acid catabolic pathways, to be incorporated into the TCA cycle as succinyl-CoA. Genes encoding the terminal reactions of branched chain amino acid degradation were not identified, although the genome encodes numerous acyl-CoA dehydrogenases with unknown specificity that could perform these functions. However, we could confidently identify the branched chain keto-acid dehydrogenase complex (BCKAD) that is critical for the degradation of leucine, isoleucine, and valine, as well as in processing the downstream degradation products of methionine and threonine. Finally, this organism carries multiple independent branched chain amino acid transport systems, as well as a polar amino acid transport system that is enriched in the Ca. Angelarcheales order **(Fig. 4b, Supplementary Fig. 6c-d, and Supplementary Table 12)**.

Angelarchaeales-1 and ammonia oxidizing Nitrososphaerales both have respiratory chains that include a complex I lacking the E, F, and G subunits for NADH binding and a duplicated subunit M that may mediate translocation of an additional proton^41^. The electron donor to the complex I may be reduced ferredoxin^41,42^. Both groups encode a four-subunit succinate/fumarate dehydrogenase, a cytochrome b-like complex III with an associated plastocyanin-like electron transfer protein, and an oxygen utilizing cytochrome c terminal oxidase complex.

Unlike the Nitrososphaerales, Angelarchaeales-1 encodes a multitude of systems for the putative utilization of ferredoxin. This includes a FixABCX electron bifurcation system that can couple the reduction of ferredoxin and quinone to the oxidation of NADH. Interestingly the FixABCX complex is co-located with the BCKAD complex in the Angelarchaeales-1 genome. This FixABCX complex may be important for converting the reducing power of NADH derived from BCKAD mediated branched chain amino acid degradation into reducing power in the form of reduced ferredoxin and quinone. Angelarchaeales-1 has 2 genes proximal to the cytochrome b-like complex III, a hdrD-like gene and a rnfB-like gene that are not found in Nitrososphaerales genomes. In rnf complexes, rnfB binds and oxidizes reduced ferredoxin, however the functions of these iron-sulfur cluster containing proteins, which are not normally found associated with complex III, are still unclear.

## DISCUSSION

In this study, we identified a novel CuMMO occurring in a largely unstudied lineage of archaea that we show is widely distributed in soils, terrestrial sediments, and even detected in deep ocean systems. The ability of a CuMMO to function as an ammonia monooxygenase requires a metabolic context that supports the acquisition of ammonia, processing of ammonia oxidation byproducts, and acquisition of energy from these reactions. Thus, the question of whether the Angelarcheales CuMMOs function as ammonia vs. methane monooxygenases can be addressed in part by comparison of the genomic context to those of ammonia monooxygenases in Nitrososphaerales. The Nitrososphaerales encode transporters for ammonia uptake. These are also present in Angelarcheales, but probably are not needed, given the ability to derive ammonia from amino acid degradation. Further, we show (i) Conservation between the protein architecture and active site residues in known archaeal amoA, amoB, and amoC ammonia monooxygenases and those found in Ca. Angelarcheales. (ii) A large inventory of cupredoxin-like BCPs including BCP subtypes (such as 2-domain and medium length 1-domain BCPs) in both Ca. Angelarcheales and Nitrososphaerales relative to sibling lineages of their respective phyla. (iii) A combination of electron transport proteins that is shared by Ca. Angelarcheales and Nitrososphaerales, consistent with similar electron transport chain functionality and energy acquisition strategies.

The presence of extensive inventories of blue copper proteins is of particular interest, as Nitrososphaerales lack a hao homologue, and BCPs are thought to perform the yet unresolved enzymatic steps that oxidize hydroxylamine to nitrite^16,17^. Work in Nitrososphaerales generally finds that while having a large number of these proteins is a clear signature of this group, the exact gene set of BCPs present in any Nitrososphaerales genome is highly variable^16^. Candidate enzymes including specific plastocyanin-like cupredoxins^19^ or nirK homologues^21,43^ have been proposed to carry out specific steps in hydroxylamine oxidation, however no individual enzyme is completely conserved across known ammonia oxidizing archaeal genomes. Thus, a variety of diverse BCPs may substitute for the missing hao activity. In Ca. Angelarcheales, we observe a similar pattern, i.e., an extensive inventory of BCPs relative to sister lineages with no individual BCP ortholog being conserved across the entire group. Instead, we observe sets of 2-domain BCPs and 1-domain medium length BCPs with divergent sequences but conserved architecture that are largely restricted to the Ca. Angelarcheales and Nitrososphaerales lineages. BCP architectures are in fact well conserved across the entire lineages, respectively. Overall, the weight of genomic evidence supports the conclusion that the Ca. Angelarcheales are a new and distinct group of ammonia-oxidizing archaea.

Ecological support for the deduction that Ca. Angelarcheales are ammonia oxidizers is their strong association with, and high abundances in, relatively aerobic grassland soils, where organisms with the capability for methanogenesis have not been detected^23,44^. These archaea also occur in sediments from above the water table and in saturated sediments that periodically receive oxygenated water^45^. In this context, it is interesting to note that the Ca. Angelarcheales can utilize amino acids as a carbon and nitrogen source without oxygen, generating energy via substrate level phosphorylation (as we detect the ability to ferment acetyl-CoA to ethanol), and oxidizing ammonia when O_2_ is available.

While we acknowledge that these data cannot represent all soils and sediments, it is striking that Ca. Angelarcheales can comprise a reasonably large fraction of communities in which they were detected. Also we find that the abundances of Ca. Angelarcheales are positively associated with both Nitrososphaerales and Nitrospirales, two groups that participate in nitrification. We note that while Ca. Angelarcheales shared significant positive associations with a number of other microbial groups, their associations with both Nitrososphaerales and Nitrospirales were some of the strongest detected. While the abundances of Angelarcheales and Nitrososphaerales are positively associated, we predict that the niches of Angelarcheales and Nitrososphaerales do not overlap, given that Angelarcheales can use ammonia that they liberate from breakdown of amino acids whereas Nitrososphaerales likely use ammonia (and urea) directly. The strong association of both Angelarcheales and Nitrososphaerales with the nitrite oxidizing Nitrospirales is consistent with the former groups producing nitrite from ammonia oxidation which is then exchanged with and oxidized by the Nitrospirales.

The identification of a lineage of archaea outside of the Nitrososphaerales involved in ammonia oxidation has important implications for biogeochemical nitrogen cycling. These archaea are unusual, as they both degrade proteins/amino acids and oxidize the product ammonia. Angelarcheales likely occur in a variety of soils and sediments where ammonia oxidation has gone undetected using primer-based methods. The new clade of CuMMOs provides an opportunity for further study of these enzymes, and for molecular clock-based analyses of their evolution relative Earth history^46^.

## METHODS

### Amo/Pmo identification, genome set selection, and dereplication

Previous genome-resolved work at the Angelo Coast Range Reserve^23^ identified 7 Thermoplasmatota genomes each containing a hit to the KOFAM Hidden Markov Model (HMM) for amoA/pmoA (K10944), below the HMM score threshold, but with a significant E-value (≤ 1E^-4^). Proteins co-localized with these divergent amoA/pmoA proteins were identified. To search for additional similar divergent amoA/pmoA proteins we aligned the divergent proteins using MAFFT v7.471 (--maxiterate 1000 --localpair) and constructed an HMM using hmmbuild in the HMMER v3.3.1 package^47^ with default parameters. This HMM was scored against all archaeal genomes in the Genome Taxonomy Database (GTDB) release r95^48^, and all archaeal MAGs in ggKbase (ggKbase.berkeley.edu) datasets as of January 13, 2020 using the hmmsearch function of the HMMER package with an HMM score threshold of 100. These genomes were phylogenetically classified using the GTDB tool kit^49^ (GTDB-Tk v1.3.0) classify workflow with default parameters. All genomes were placed within the RBG-16-68-12 order (hereafter Ca. Angelarcheales) of the GTDB taxonomy (**Supplementary Table 1**).

To produce a full genome reference set for our analyses we added an additional 31 genomes that did not have any hit to our custom amoA/pmoA HMM but fell within the Ca. Angelarcheales order (**Supplementary Table 1**). These genomes came both from Angelo Coast Range Reserve assemblies (n = 15 genomes) and from the Ca. Angelarcheales order in GTDB (n = 16 genomes). We also added 719 reference genomes from GTDB derived from the archaeal phyla Thermoplasmatota (n = 338 genomes) and Thermoproteota (n = 371). Thermoproteota genomes were included for comparison of functional differences within and between phyla of known (Nitrososphaerales) and putative (Ca. Angelarcheales) ammonia oxidizing archaea.

The full genomes set was de-replicated at the species level (Average Nucleotide Identity ≥ 95 %) using dRep v3.0.1^50^ with the following parameters: -p 16 -comp 10 -ms 10000 -sa 0.95. The best genome from each species cluster was chosen as a representative genome by dRep. Species representatives were required to have ≥ 60 % completeness and ≤ 10 % contamination as estimated by checkM^51^. If no genome within a species cluster met these criteria the cluster was discarded. All genome information can be found in **Supplementary Tables 2 and 4**.

### Amo/Pmo protein identification, alignment, analysis, and phylogenetic reconstruction

The amoA/pmoA protein was used as an anchor sequence to manually annotate putative amoB/pmoB, amoC/pmoC, and amoX proteins present on each contig across the Ca. Angelarcheales genomes (**Supplementary Table 2**). Manual annotation used a combination of protein order relative to the amoA/pmoA sequence, predicted protein length (compared to known amo/pmo proteins), and best blast hits vs. the NCBI nr database.

To identify amoB/pmoB and amoC/pmoC in Ca. Angelarcheales genomes where no amoA/pmoA was identified, and to develop a method to rapidly identify all novel amo/pmo complex proteins in future work we aligned each set of amo/pmo proteins following manual annotation using MAFFT v7.471 (--maxiterate 1000 --localpair) and constructed HMMs for each using hmmbuild in the HMMER v3.3.1 package with default parameters. These were scored against all proteins in the full set of 228 redundant Ca. Angelarcheales genomes (**Supplementary Table 1**) using the hmmsearch function of the HMMER v3.3.1 package with an HMM score threshold of 100.

Alignments of putative amo/pmo sequences with known references were constructed by merging amo/pmo sequences from Ca. Angelarcheales were with reference sequences from Khadka, et al.^1^ and those from an additional set of 22 ammonia oxidizing archaeal reference genomes (**Supplementary Table 3**). This resulted in protein sets for amoA/pmoA, amoB/pmoB, and amoC/pmoC that contained 112, 114, and 110 sequences respectively. Sequence sets were aligned using MAFFT v7.471 with the following parameters: --maxiterate 1000 --localpair --reorder --thread 12. Mean amino acid identity between Ca. Angelarcheales, bacterial, and archeal reference sequences were calculated in Geneious Prime v2020.2.4 from a pairwise sequence identity matrix. Conserved residues for methane and ammonia monooxygenases were referenced from Wang, et al.^29^ and identified in each alignment through manual inspection.

Maximum likelihood phylogenetic trees were constructed for each individual amo/pmo subunit alignment using IQ-TREE v1.6.12^52^ with the following options: -m MFP -bb 1000 -alrt 1000 -nt 12. The empirically selected evolutionary rate model for all amo/pmo sequence sets was LG+F+I+G4 based on Bayesian information criteria (BIC). Branch support was estimated using ultrafast bootstrapping with 1000 bootstrap replicates. Individual amo/pmo protein trees can be found in **Supplementary Figure 3**. For the combined amoABC/pmoABC tree, sequences from the same organism were concatenated in Geneious Prime v2020.2.4 and only retained if at least 2 of the proteins were present, resulting in 112 total concatenated sequences in the alignment. A Maximum likelihood phylogenetic tree was constructed for concatenated sequences using IQ-TREE v1.6.12 with the following options: -m MFP -bb 1000 -alrt 1000 -nt 12. The empirically selected evolutionary rate model for the concatenated amoABC/pmoABC tree was LG+F+I+G4 based on BIC. Branch support was estimated using ultrafast bootstrapping with 1000 bootstrap replicates.

### Assessment of AmoA/PmoA primer complementarity

Primer sequences used to amplify and quantify both archaeal (GenAOAF: 5’-ATA GAG CCT CAA GTA GGA AAG TTC TA-3’ ; GenAOAR: 5’-CCA AGC GGC CAT CCA GCT GTA TGT CC-3’) and bacterial (amoA-1Fmod: 5’-CTG GGG TTT CTA CTG GTG GTC-3’ ; GenAOBR: 5’-GCA GTG ATC ATC CAG TTG CG-3’) amoA/pmoA sequences using PCR from environmental samples were obtained from Meinhardt, et al.^28^. Sequence complementarity was assessed using the map primers function of Geneious Prime v2020.2.4. Primers were tested against nucleotide sequences of amoA genes from Nitrososphaerales genomes and amoA/pmoA genes from Ca. Angelarcheales genomes as detailed in **Supplementary Table 3**. A match required both the forward and reverse primers to bind to the sequence while allowing up to 7 mismatched bases in each primer. Full data obtained for the number of mismatches and estimated product sizes for each primer pair and template sequence are available in **Supplementary Table 3**.

### Genome taxonomy and phylogenetic reconstruction

Initial taxonomic placement for the 645 non-redundant genomes used in this study was performed using the GTDB-Tk^49^ classify workflow with the following parameters: classify_wf -x fasta --cpus 48. All GTDB-Tk based taxonomic classification is available in **Supplementary Table 4**. A concatenated marker gene phylogenetic tree for all genomes classified within the archaeal phylum Thermoplasmatota was constructed by combining the 34 Ca. Angelarcheales genomes with 302 de-replicated reference genomes from this phylum spanning all known orders. The *Archaeoglobus fulgidus* (GCF_000008665.1) genome was also included to be used as an outgroup for tree rooting. GToTree v1.5.22^30^ was used to identify and extract a set of 76 phylogenetically informative single copy archaeal marker genes (SCGs), defined in Lee, et al.^30^, from each genome using the following parameters: -H Archaea.hmm -j 8 -d. Genomes where < 50% (38 genes) of the targeted marker genes could not be identified were removed from the analysis (n = 10 genomes) retaining a total of 327 genomes in the final tree. SCG sequence sets were then individually aligned with Muscle v3.8.31^53^ and alignments were trimmed with Trimal v1.4^54^. All alignments were then concatenated, and an SCG alignment partition table was produced by GToTree so evolutionary substitution rate models could be estimated for each SCG independently during phylogenetic tree construction. A maximum likelihood phylogenetic tree was constructed with IQ-TREE v1.6.12^52^ with the following options: -spp Partitions.txt -m MFP -bb 1000 -alrt 1000 -nt 48. Evolutionary rate models were empirically estimated for each marker gene independently and selected based on BIC. Branch support was estimated using ultrafast bootstrapping with 1000 bootstrap replicates. Phylogenetic trees were rooted using *Archaeoglobus fulgidus* as an outgroup, annotated, and displayed using iTOL v6. For the full tree see **Supplementary Figure 4**.

### RpL6 marker abundance and association analysis

There are 6 study sites where at least 1 Ca. Angelarcheales genome was reconstructed from shotgun metagenome data. We used all 185 shotgun metagenomic samples collected from these sites, regardless of whether a Ca. Angelarcheales genome was recovered from a sample, to estimate the relative abundance of the Ca. Angelarcheales order across these locations (**Supplementary Table 5)**. Relative abundance of all bacterial and archaeal order level taxonomic groups was quantified using marker gene taxonomic placement and quantification of the phylogenetically informative SCG ribosomal protein L6 (rpL6). RpL6 marker gene profiling and quantification was performed using GraftM v0.13.1^55^. Briefly, an rpL6 graftM database was constructed using the ribosomal L6 protein sequences from all archaeal and bacterial genomes (provided by GTDB using TIGR03653 and TIGR03654, respectively) in GTDB v95. A GraftM package was then created using this set of sequences and the rpL6 Pfam HMM (PF00347.24) sensitive for both bacterial and archaeal variants using the command: graftM create --sequences L6.faa --taxonomy taxonomy.tsv --hmm PF00347.hmm. GraftM was then used to call genes, identify rpL6 sequences, phylogenetically place sequences, and quantify read counts of rpL6 sequences identified in each sample using the following options: graftM graft --forward [forward reads] --reverse [reverse reads] --graftm_package rpL6_gpkg --threads 48. Raw counts in each sample were aggregated to the taxonomic rank of order using a custom R script (https://github.com/SDmetagenomics/AMO_Archaea_2021). Counts of rpL6 sequences that could only be resolved to taxonomic ranks higher than order level were retained in their original form. All aggregated count data is available in **Supplementary Table 6**. Relative abundances of the Ca. Angelarcheales and Nitrososphaerales orders in each sample were calculated as the fraction of total reads in a sample that were associated with these order ranks. Plotting was performed in R using the ggplot2 package^56^.

Association analysis between the abundances of each taxonomic group across samples (**Supplementary Table 7**) was conducted using the propr package^57^ in R. Proportionality was chosen as an association measure over correlation as it is better suited for compositional data such as sequencing read counts^58^. Briefly, taxonomic groups were filtered such that only those with ≥ 5 counts in at least 33 % of samples (n = 61 samples) were retained. This filtering was performed to remove taxonomic groups with extremely low counts from association comparisons as to avoid spurious associations. Following filtering 90.5 % of the original count data was retained in each sample on average. Subsequently the rho proportionality metric was calculated for centered log-ratio normalized counts between all pairs of taxonomic groups using the following function: propr(count_matrix, metric = “rho”, p = 1000). The magnitude of proportionality that represented a significant association (FDR < 0.001) was calculated through a permutation based procedure implemented using the updateCutoffs function within the propr package as follows: updateCutoffs(L6_rho, cutoff = seq(from = 0.05, to = 0.40,by = 0.01), ncores = 12). It was determined that all rho values ≥ 0.25 were statistically significant with a FDR < 0.001. Additionally, only positive rho association values were considered in the final analysis, as negative associations in compositional data are considered less reliable, even after compositional correction measures are applied^57^. All significant proportionality values are available in **Supplementary Table 7**. The abundance values in the plotted pairwise comparisons between Ca. Angelarcheales, Nitrososphaerales, and Nitrospirales use the centered log-ratio of counts in each sample as calculated by the propr package. All plotting was performed using the ggplot2 package in R.

### Genome annotation

For the 645 genomes passing completeness and contamination quality criteria, annotation was performed as follows. Genes and protein sequences were predicted using prodigal v2.3.6^59^ using the following options: prodigal -i [genome contigs] -a [proteins sequences out] -m -p meta. KEGG KO annotations were predicted using KofamScan^60^ using HMM models from release r02_18_2020 with the following options: exec_annotation -p [hmm profiles] -k [hmm cutoffs] --cpu 48 --tmp-dir [temp dir] -o [output folder] [protein file]. As multiple KEGG HMMs can match to the same protein with scores exceeding their score cutoff thresholds, the HMM with the lowest E-value had its annotation transferred to the protein. If a protein did not match any KEGG HMM above the HMM cutoff threshold then the lowest E-value annotation was transferred. An E-value cutoff of < 1e-10 was applied, above which no annotations were transferred, and genes were not assigned to a KO. Archeal COG (arCOGs) annotations were predicted for all proteins using HMMs from EggNOG v5^61^ using the following options: hmmsearch --tblout [arCOG hit table] -E 0.0001 --cpu 10 All_arCOG.hmm All_Proteins.faa. We searched for XoxF/mxaF-like pqq-binding methanol dehydrogenases in genomes using the hmmsearch function of the HMMER package and a custom HMM from Anantharaman, K. et al.^26^ with an HMM score threshold cutoff of ≥ 166 using the following options: hmmsearch --tblout [xoxF_mxaF hits] -T 166 --cpu 12 methanol_dehydrogenase_pqq_xoxF_mxaF.hmm All_Proteins.faa. Protease counts in each genome were determined via the METABOLIC pipeline (https://github.com/AnantharamanLab/METABOLIC) implemented using the following options: perl METABOLIC-G.pl -t 48 -m-cutoff 0.75 -in [input protein files] -kofam-db full -o [output annotations]. Custom R code for parsing and plotting METABOLIC outputs used for peptidase quantification can be found in our github repository. Pfam domain annotations were predicted by searching all proteins against the PfamA database release r32^62^ using the hmmsearch function of the HMMER package with the following options: hmmsearch --domtblout [pfam domain table] --cut_ga --cpu 10 Pfam-A.hmm All_Proteins.faa. Overlapping pfam domain matches to the same protein were resolved, and domain boundaries were established, using the cath-resolve-hits function of the cath-tools package with default parameters (https://github.com/UCLOrengoGroup/cath-tools). All annotations were aggregated into a final table using a custom R script which is available in our github repository. Also the complete gene level annotation table for all 645 genomes is available at FigShare (https://figshare.com/projects/AMO_Archaea/112599).

For functional annotations of genes and pathways across all Ca. Angelarcheales genomes, a subset of 190 specific target functions were analyzed (**Fig. 2b and Supplementary Figure 6a)**. Pathways and protein complexes were subsequently grouped into 77 “gene groups” that represent the capability to perform a pathway or metabolic function. Each gene group had specific criteria that were required (i.e. a critical protein needed to be present) for a positive detection. All targets, gene group organization, and criteria for detection are available in **Supplementary Table 8**. For annotation and quantification of amino acid and peptide transport systems across all 645 genomes in our analysis a set of 38 KEGG KOs representing individual transporter proteins or transporter subunits were used as the search criteria (**Supplementary Figure 6c-d and Supplementary Table 14**). For functional annotation of the Angelarcheales-1 genome and metabolic map reconstruction (**Fig. 5**), a subset of 233 specific target functions were analyzed that highlighted central carbon metabolism, the degradation of amino acids, and electron transport/energy generation (**Supplementary Table 12)**. Pathways and protein complexes were subsequently grouped into 115 “reaction groups” that represent the capability to perform a pathway or metabolic function. Each reaction group had specific criteria that were required (i.e. a critical protein needed to be present) for a positive detection. All targets, reaction group organization, genes associated with reactions, and criteria for detection are available in **Supplementary Table 12**.

### Protein clustering

Clustering of all proteins predicted in the 645 genomes passing completeness and contamination quality criteria was accomplished as described in Méheust, et al.^33^. Code for this pipeline is available at: https://github.com/raphael-upmc/proteinClusteringPipeline. Briefly, all 1,103,913 proteins were first clustered into subfamilies (subfams) using the subfams.py script of the pipeline (which implements mmseqs2^63^ clustering) using the following options: subfamilies.py -- output-directory [clustering dir] --cpu 48 --coverage 0.8 All_Proteins.faa. All proteins within a subfam must align with bidirectional coverage of at least 80 % (--cov-mode 0 in mmseqs2), and alignments must have an E-value < 1e-4. Proteins which did not cluster into a subfam of at least 2 proteins were discarded from this analysis leaving a total of 964,644 (87.4 %) proteins with subfamily assignments. A total of 76,216 protein subfam clusters were formed. Subsequently proteins within each subfam were aligned, HMMs were constructed from these alignments, and all-v-all HMM scoring was conducted using the hhblits.py script (which implements the hhblits function within the hhsuite v3.0^64^ software package) using default options. For HMM-HMM scoring any local alignments between HMMs of different subfams had to have an hhblits probability score of ≥ 95 % to be retained for downstream analysis. Finally, family groupings of subfams (fams) were formed by applying the Markov clustering algorithm (MCL)^65^ to the network of all HMM-HMM connections using the runningMclClustering.py script of the pipeline using the following options: runningMclClustering.py --force --min-size 2 --cpu 4 --fasta config.json. This resulted in the formation of 19,828 protein fam clusters.

### BCP identification, quantification, and phylogenetics

Cupredoxin-like blue copper proteins (BCPs) were initially identified in our dataset by selecting all proteins that carried one of the following Pfam domains: Copper-bind, COX2, COX_ARM, Cu-oxidase, Cu-oxidase_2, Cu-oxidase_3, Cu_bind_like, Cupredoxin_1, CzcE, DP-EP, Ephrin, hGDE_N, PAD_N, PixA, SoxE. These domains are all members of the Pfam CU_oxidase clan (CL0026). Due to the large sequence divergence in cupredoxin-like domains (as evidenced by the number of pfam models required to appropriately capture their diversity), we posited that many divergent domain sequences would be missed by direct Pfam annotation. Alternatively, more sensitive local HMM-HMM comparison at the subfam level would cluster the majority of BCP domain containing proteins into a single fam cluster. Thus, we quantified the number of BCP domain containing proteins present in all fams. Given that 90.2% of proteins with annotated BCP domains were members of fam00018, we used fam00018 to represent BCP domain containing proteins in our dataset. Comparison of fam00018 protein counts in genomes was carried out independently for the archaeal phyla Thermoplasmatota and Thermoproteota between all archeal orders within these phyla that contained ≥ 3 genomes. Global significant differences across all orders within a phylum was first tested using the Kruskal-Wallis rank sum test implemented as the kruskal.test() function in R (α ≤ 0.05). Pairwise significant differences between orders within a phylum was then tested using the Wilcoxon rank sum test implemented as the pairwise.wilcox.test() function in R. P-values from tests were corrected for multiple comparisons using false discovery rate (FDR) with a value of FDR ≤ 0.05 being considered significant. All aggregation, quantification, statistical testing, and plotting of BCP protein count data was performed using a custom R script available in our github repository.

Manual subfamily level annotation and domain architecture analysis was performed on 100 fam00018 subfams that contained at least 5 proteins (85.2% of all fam00018 proteins). The proteins in each subfam were re-aligned with MAFFT v7.471 using the following options: mafft -- maxiterate 1000 --localpair --reorder --thread 12 [proteins in] > [alignment out]. HMMs were built for each alignment using hhsuite v3.0, and HMM-HMM scoring was performed against the PfamA database release r32 using hhsearch with the following options: hhsearch -i [input hmm] -o [output table] -d [pfam hmm database] -p 50 -E 0.001 -z 1 -Z 32000 -b 0 -B 0 -n 1 -cpu 18. Domain matches with a probability score of ≥ 95 % were retained, and Pfam domains overlapping the same region on target subfam HMMs were resolved with the cath-resolve-hits function of the cath-tools package using default parameters. Each of the 100 subfams were then manually annotated and placed into 1 of 6 broad classification groups based on the subfam level Pfam domain architectures and the types of KEGG and arCOG annotations assigned to individual proteins within the subfam. Annotation groups were defined as follows: COX2 (Contains COX2 domain and KEGG or arCOG annotations indicate > 40% of proteins in cluster are terminal oxidase subunits), 1D_Cu_Small (one Pfam BCP domain, mean protein length < 250 amino acids), 1D_Cu_Med (one Pfam BCP domain, mean protein length 250 - 400 amino acids), 1D_Cu_Large (one Pfam BCP domain, mean protein length > 400 amino acids), 2D_Cu (two Pfam BCP domains), 3D_Cu (three Pfam BCP domains). Due to its presence in Angelarcheales-1 and its proximity to the coxAB locus subfam00588 was also manually annotated as above despite having < 5 proteins. Assignment of copper site types (e.g. Type 1 Copper) was conducted by manual inspection of subfamily alignments and referenced from Gräff, et al.^35^. All subfams within fam00018 and their associated annotations are available in **Supplementary Table 9**.

Phylogenetic tree construction for manually annotated subfams with > 1 BCP domain was undertaken as follows: All proteins from subfams in the 2D_Cu and 3D_Cu manual annotation groups (n = 349 proteins) were combined with reference laccase and nitrite reductase (nirK) sequences from Decleyre, et al.^66^, Kobayashi, et al.^36^, and Nakamura, et al.^67^ (n = 90 proteins). Proteins were aligned with MAFFT v7.471 using the following options: mafft --maxiterate 1000 -- genafpair --reorder --thread 12 [proteins in] > [alignment out]. A maximum likelihood phylogenetic tree was constructed using IQ-TREE v1.6.12 with the following options: -m MFP -bb 1000 -alrt 1000 -nt 12. The empirically selected evolutionary rate model for the tree was WAG+F+R7 based on BIC. Branch support was estimated using ultrafast bootstrapping with 1000 bootstrap replicates. Tree was annotated, and displayed using iTOL v6. Tree clades were manually defined based on positioning of reference sequences within clusters and copper site types present in sequences defined in Nakamura, et al.^67^.

### Gene co-occurrence analysis and locus plotting

Contigs containing genes for the amoCAXB cluster, the coxAB oxygen utilizing terminal oxidase cluster, and the complex III like cytochrome b gene cluster were identified in Ca. Angelarcheales genomes as follows: amoCAXB contigs were identified as any contig encoding an amo/pmo subunit (as described above); coxAB containing contigs were identified as any contig encoding a gene that matched to the KEGG HMM for K02274 (coxA); and complex III like gene cluster containing contigs were identified as any counting encoding a gene that matched the arCOG HMM for arCOG01721 (cytochrome b of bc complex). All loci were extracted from our master annotation table using a custom R script available in our github repository, and loci were displayed using the gggenes package in R (https://github.com/wilkox/gggenes). For ease of viewing amoCAXB and coxAB containing contigs were truncated to display 10 genes on either side of the gene cluster of interest. Complex III containing contigs were truncated to display 22 genes on either side of the gene cluster as to allow inclusion of other proximal respiratory complexes.

### KEGG Ortholog (KO) enrichment analysis

Detection of enriched KO terms between archaeal orders was carried out using a custom R script available in our github repository. Archaeal orders from the phyla Thermoplasmatota and Thermoproteota that contained < 3 genomes were excluded from the analysis. For the remainder of genomes, a table was generated giving the count of all observed KOs (n = 5,929 total unique KO terms) in every genome. KOs that occurred less than 10 times across all genomes were filtered from the analysis. The filtered KO count table along with the taxonomic order level groupings of genomes were used as the input for indicator species analysis implemented as the multipatt() function in the R indicspecies package^68^ using the following options: multipatt(x = [KO count matrix], cluster = [genome taxonomic assignment], func = “IndVal.g”, max.order = 2, restcomb = c(1:25, 262), control = how(nperm = 9999)). We analyzed the enrichment of KO term frequency in all orders individually as well as a grouping consisting of the combined genomes from orders Ca. Angelarcheales and Nitrosospherales. This allows for the identification of enriched functions that are shared by these two groups relative to all other lineages and to each other individually. Statistical significance of enriched KO frequency was estimated by permutation using 9,999 permuted groupings of the genomes. P-values were corrected for multiple testing using FDR with a value of FDR ≤ 0.05 being considered significant. Additionally only KOs with an indicator value ≥ 0.4 were retained for downstream analysis. KOs enriched in Nitrosospherales (Gp 14), Ca. Angelarcheales (Gp 17), or shared by both orders (Gp 26) were displayed as a heatmap using the superheat package in R. For a list of all significant KOs and their functions see **Supplementary Table 10**. Functional category assignments for KOs were derived from KEGG orthology group hierarchies and manually curated, and quantified in R. The full list of KO to functional category assignments is available in our github repository.

## DATA AVAILABILITY

Genomic data, including assembled genomes and raw sequencing reads, are available under the following NCBI BioProject accession numbers: PRJNA449266 and PRJNA288027. Ca. Angelarcheales genomes from the project can also be found at ggKbase (https://ggkbase.berkeley.edu/AMO_Archaea). Large datasets including an ORF-to-Genome table, all predicted proteins, and the full gene level annotation table are available at figshare (https://figshare.com/projects/AMO_Archaea/112599).

## CODE AVAILABILITY

Code use for the analysis in this report are available at the following github repository: https://github.com/SDmetagenomics/AMO_Archaea_2021

## REPORTING SUMMARY

Further information on research design is available in the Nature Research Reporting Summary linked to this article.

## ACKNOWLEDGEMENTS

The authors thank Ka Ki (Lily) Law and Shufei Lei for data support and Alex Thomas for helpful discussion, and the Joint Genome Institute for a subset of the sequencing. We acknowledge Dong, et al. (2019) whose public data we used in this study. This work was facilitated by samples previously collected at the University of California Natural Reserve system Angelo Coast Range Reserve (10.21973/N3R94R). Funding was provided by the Office of Science, Office of Biological and Environmental Research, of the US Department of Energy (grant DOE-SC10010566), and by m-CAFEs (Microbial Community Analysis & Functional Evaluation in Soils), a project led by Lawrence Berkeley National Laboratory supported by the U.S. Department of Energy, Office of Science, Office of Biological & Environmental Research under contract number DE-AC02-05CH11231.

## AUTHOR CONTRIBUTIONS

SD, AL, and KHW collected samples. SD, AL and JFB designed the study. SD, AL, AS, and JFB assembled and curated genomes. SD, AL, ACC, and JFB performed genome annotation, bioinformatic and statistical analysis. SD and ACC developed code. SD, PBMC, and JFB interpreted metabolic annotation data and curated metabolic pathways. SD, AL, ACC, and JFB wrote the manuscript. All others read and approved the content.

## CORRESPONDING AUTHORS

Correspondence should be addressed to Spencer Diamond and Jillian F. Banfield.

## REFERENCES

1. Khadka, R. et al. Evolutionary History of Copper Membrane Monooxygenases. Frontiers in microbiology 9, W132–13 (2018).

2. Coleman, N. V. et al. Hydrocarbon monooxygenase in Mycobacterium: recombinant expression of a member of the ammonia monooxygenase superfamily. The ISME Journal 6, 171–182 (2011).

3. Lehtovirta-Morley, L. E. Ammonia oxidation: Ecology, physiology, biochemistry and why they must all come together. FEMS Microbiology Letters 365, 28 (2018).

4. Stahl, D. A. & Torre, J. R. de la. Physiology and Diversity of Ammonia-Oxidizing Archaea. Annual Review of Microbiology 66, 83–101 (2012).

5. Mancinelli, R. L. The Regulation of Methane Oxidation in Soil. Annu Rev Microbiol 49, 581–605 (1995).

6. Wu, L. et al. A Critical Review on Nitrous Oxide Production by Ammonia-Oxidizing Archaea. Environmental Science & Technology 54, 9175–9190 (2020).

7. Monteiro, M., Séneca, J. & Magalhães, C. The history of aerobic ammonia oxidizers: from the first discoveries to today. Journal of Microbiology 52, 537–547 (2014).

8. Alves, R. J. E., Minh, B. Q., Urich, T., Haeseler, A. von & Schleper, C. Unifying the global phylogeny and environmental distribution of ammonia-oxidising archaea based on amoA genes. Nature communications 9, 66–17 (2018).

9. Knief, C. Diversity and Habitat Preferences of Cultivated and Uncultivated Aerobic Methanotrophic Bacteria Evaluated Based on pmoA as Molecular Marker. Frontiers in microbiology 6, 457–38 (2015).

10. Hatzenpichler, R. Diversity, Physiology, and Niche Differentiation of Ammonia-Oxidizing Archaea. Applied and Environmental Microbiology 78, 7501–7510 (2012).

11. Leininger, S. et al. Archaea predominate among ammonia-oxidizing prokaryotes in soils. Nature 442, 806–809 (2006).

12. Liu, S. et al. Ammonia-oxidizing archaea have better adaptability in oxygenated/hypoxic alternant conditions compared to ammonia-oxidizing bacteria. Applied Microbiology and Biotechnology 99, 8587–8596 (2015).

13. Tolar, B. B. et al. Integrated structural biology and molecular ecology of N-cycling enzymes from ammonia-oxidizing archaea. Environmental Microbiology Reports 9, 484–491 (2017).

14. Bédard, C. & Knowles, R. Physiology, biochemistry, and specific inhibitors of CH4, NH4+, and CO oxidation by methanotrophs and nitrifiers. Microbiol Rev 53, 68–84 (1989).

15. Burrows, K. J., Cornish, A., Scott, D. & Higgins, I. J. Substrate Specificities of the Soluble and Particulate Methane Mono-oxygenases of Methylosinus trichosporium OB3b. Microbiology+ 130, 3327–3333 (1984).

16. Qin, W. et al. Alternative strategies of nutrient acquisition and energy conservation map to the biogeography of marine ammonia-oxidizing archaea. The ISME Journal 409, 507–15 (2020).

17. Lancaster, K. M., Caranto, J. D., Majer, S. H. & Smith, M. A. Alternative Bioenergy: Updates to and Challenges in Nitrification Metalloenzymology. Joule 2, 421–441 (2018).

18. Vajrala, N. et al. Hydroxylamine as an intermediate in ammonia oxidation by globally abundant marine archaea. Proceedings of the National Academy of Sciences 110, 1006–1011 (2013).

19. Hosseinzadeh, P. et al. A Purple Cupredoxin from Nitrosopumilus maritimus Containing a Mononuclear Type 1 Copper Center with an Open Binding Site. J Am Chem Soc 138, 6324–6327 (2016).

20. Stein, L. Y. Insights into the physiology of ammonia-oxidizing microorganisms. Current Opinion in Chemical Biology 49, 9–15 (2019).

21. Carini, P., Dupont, C. L. & Santoro, A. E. Patterns of thaumarchaeal gene expression in culture and diverse marine environments. Environmental Microbiology 20, 2112–2124 (2018).

22. Pester, M. et al. amoA-based consensus phylogeny of ammonia-oxidizing archaea and deep sequencing of amoA genes from soils of four different geographic regions. Environmental Microbiology 14, 525–539 (2012).

23. Diamond, S. et al. Mediterranean grassland soil C-N compound turnover is dependent on rainfall and depth, and is mediated by genomically divergent microorganisms. Nature Microbiology 4, 1356–1367 (2019).

24. Sharrar, A. M. et al. Bacterial Secondary Metabolite Biosynthetic Potential in Soil Varies with Phylum, Depth, and Vegetation Type. Mbio 11, e00416–20 (2020).

25. Lavy, A. et al. Microbial communities across a hillslope-riparian transect shaped by proximity to the stream, groundwater table, and weathered bedrock. Ecol Evol 9, 6869–6900 (2019).

26. Anantharaman, K. et al. Thousands of microbial genomes shed light on interconnected biogeochemical processes in an aquifer system. Nature communications 7, 13219 (2016).

27. Dong, X. et al. Metabolic potential of uncultured bacteria and archaea associated with petroleum seepage in deep-sea sediments. Nat Commun 10, 1816 (2019).

28. Meinhardt, K. A. et al. Evaluation of revised polymerase chain reaction primers for more inclusive quantification of ammonia-oxidizing archaea and bacteria. Env Microbiol Rep 7, 354–363 (2015).

29. Wang, V. C. C. et al. Alkane Oxidation: Methane Monooxygenases, Related Enzymes, and Their Biomimetics. Chemical Reviews 117, 8574–8621 (2017).

30. Lee, M. D. GToTree: a user-friendly workflow for phylogenomics. Bioinformatics 35, btz188 (2019).

31. Lawton, T. J., Ham, J., Sun, T. & Rosenzweig, A. C. Structural conservation of the B subunit in the ammonia monooxygenase/particulate methane monooxygenase superfamily. Proteins 82, 2263–2267 (2014).

32. Bowers, R. M. et al. Minimum information about a single amplified genome (MISAG) and a metagenome-assembled genome (MIMAG) of bacteria and archaea. Nat Biotechnol 35, 725–731 (2017).

33. Méheust, R., Burstein, D., Castelle, C. J. & Banfield, J. F. The distinction of CPR bacteria from other bacteria based on protein family content. Nat Commun 10, 4173 (2019).

34. Komori, H. & Higuchi, Y. Structure and molecular evolution of multicopper blue proteins. Biomol Concepts 1, 31–40 (2010).

35. Gräff, M., Buchholz, P. C. F., Roes-Hill, M. L. & Pleiss, J. Multicopper oxidases: modular structure, sequence space, and evolutionary relationships. Proteins Struct Funct Bioinform 88, 1329–1339 (2020).

36. Kobayashi, S. et al. Nitric Oxide Production from Nitrite Reduction and Hydroxylamine Oxidation by Copper-containing Dissimilatory Nitrite Reductase (NirK) from the Aerobic Ammonia-oxidizing Archaeon, Nitrososphaera viennensis. Microbes Environment 33, ME18058 (2018).

37. Reyes, C. et al. Genome wide transcriptomic analysis of the soil ammonia oxidizing archaeon Nitrososphaera viennensis upon exposure to copper limitation. The ISME Journal 1– 16 (2020) doi:10.1038/s41396-020-0715-2.

38. Kitzinger, K. et al. Cyanate and urea are substrates for nitrification by Thaumarchaeota in the marine environment. Nature Microbiology 1–12 (2019) doi:10.1038/s41564-018-0316-2.

39. Bräsen, C., Esser, D., Rauch, B. & Siebers, B. Carbohydrate Metabolism in Archaea: Current Insights into Unusual Enzymes and Pathways and Their Regulation. Microbiol Mol Biol R 78, 89–175 (2014).

40. Tjaden, B., Plagens, A., Dörr, C., Siebers, B. & Hensel, R. Phosphoenolpyruvate synthetase and pyruvate, phosphate dikinase of Thermoproteus tenax: key pieces in the puzzle of archaeal carbohydrate metabolism. Mol Microbiol 60, 287–298 (2006).

41. Chadwick, G. L., Hemp, J., Fischer, W. W. & Orphan, V. J. Convergent evolution of unusual complex I homologs with increased proton pumping capacity: energetic and ecological implications. Isme J 12, 2668–2680 (2018).

42. Welte, C. & Deppenmeier, U. Membrane-Bound Electron Transport in Methanosaeta thermophila. J Bacteriol 193, 2868–2870 (2011).

43. Bartossek, R., Nicol, G. W., Lanzen, A., Klenk, H.-P. & Schleper, C. Homologues of nitrite reductases in ammonia-oxidizing archaea: diversity and genomic context. Environmental Microbiology 12, 1075–1088 (2010).

44. Butterfield, C. N. et al. Proteogenomic analyses indicate bacterial methylotrophy and archaeal heterotrophy are prevalent below the grass root zone. PeerJ 4, e2687–28 (2016).

45. Yabusaki, S. B. et al. Water Table Dynamics and Biogeochemical Cycling in a Shallow, Variably-Saturated Floodplain. Environ Sci Technol 51, 3307–3317 (2017).

46. Ward, L. M., Johnston, D. T. & Shih, P. M. Phanerozoic radiation of ammonia oxidizing bacteria. Sci Rep-uk 11, 2070 (2021).

47. Eddy, S. R. Accelerated Profile HMM Searches. PLoS Computational Biology 7, e1002195–16 (2011).

48. Parks, D. H. et al. A complete domain-to-species taxonomy for Bacteria and Archaea. Nature Biotechnology 1–12 (2020) doi:10.1038/s41587-020-0501-8.

49. Chaumeil, P.-A., Mussig, A. J., Hugenholtz, P. & Parks, D. H. GTDB-Tk: a toolkit to classify genomes with the Genome Taxonomy Database. Bioinformatics 1–3 (2019) doi:10.1093/bioinformatics/btz848.

50. Olm, M. R., Brown, C. T., Brooks, B. & Banfield, J. F. dRep: a tool for fast and accurate genomic comparisons that enables improved genome recovery from metagenomes through dereplication. 11, 2864–2868 (2017).

51. Parks, D. H., Imelfort, M., Skennerton, C. T., Hugenholtz, P. & Tyson, G. W. CheckM: assessing the quality of microbial genomes recovered from isolates, single cells, and metagenomes. Genome Research 25, 1043–1055 (2015).

52. Nguyen, L.-T., Schmidt, H. A., Haeseler, A. von & Minh, B. Q. IQ-TREE: A Fast and Effective Stochastic Algorithm for Estimating Maximum-Likelihood Phylogenies. Molecular biology and evolution 32, 268–274 (2014).

53. Edgar, R. C. MUSCLE: multiple sequence alignment with high accuracy and high throughput. Nucleic acids research 32, 1792–1797 (2004).

54. Capella-Gutiérrez, S., Silla-Martínez, J. M. & Gabaldón, T. trimAl: a tool for automated alignment trimming in large-scale phylogenetic analyses. Bioinformatics 25, 1972–1973 (2009).

55. Boyd, J. A., Woodcroft, B. J. & Tyson, G. W. GraftM: a tool for scalable, phylogenetically informed classification of genes within metagenomes. Nucleic Acids Res 46, gky174. (2018).

56. Wickham, H. ggplot2: Elegant Graphics for Data Analysis. (Springer Science & Business Media, 2009). doi:10.1007/978-0-387-98141-3.

57. Quinn, T. P., Richardson, M. F., Lovell, D. & Crowley, T. M. propr: An R-package for Identifying Proportionally Abundant Features Using Compositional Data Analysis. Sci Rep-uk 7, 16252 (2017).

58. Quinn, T. P. et al. A field guide for the compositional analysis of any-omics data. Gigascience 8, (2019).

59. Hyatt, D. et al. Prodigal: prokaryotic gene recognition and translation initiation site identification. BMC Bioinformatics 11, 119–11 (2010).

60. Aramaki, T. et al. KofamKOALA: KEGG ortholog assignment based on profile HMM and adaptive score threshold. Bioinformatics 36, 2251–2252 (2019).

61. Huerta-Cepas, J. et al. eggNOG 5.0: a hierarchical, functionally and phylogenetically annotated orthology resource based on 5090 organisms and 2502 viruses. Nucleic Acids Res 47, gky1085 (2018).

62. Mistry, J. et al. Pfam: The protein families database in 2021. Nucleic Acids Res 49, gkaa913. (2020).

63. Steinegger, M. & Söding, J. MMseqs2 enables sensitive protein sequence searching for the analysis of massive data sets. Nat Biotechnol 35, 1026–1028 (2017).

64. Steinegger, M. et al. HH-suite3 for fast remote homology detection and deep protein annotation. Bmc Bioinformatics 20, 473 (2019).

65. Enright, A. J., Dongen, S. V. & Ouzounis, C. A. An efficient algorithm for large-scale detection of protein families. Nucleic Acids Res 30, 1575–1584 (2002).

66. Helen, D., Kim, H., Tytgat, B. & Anne, W. Highly diverse nirK genes comprise two major clades that harbour ammonium-producing denitrifiers. BMC Genomics 17, 1–13 (2016).

67. Nakamura, K. & Go, N. Function and molecular evolution of multicopper blue proteins. Cellular and Molecular Life Sciences 62, 2050–2066 (2005).

68. Cáceres, M. D. & Legendre, P. Associations between species and groups of sites: indices and statistical inference. Ecology 90, 3566–3574 (2009).

