## Supplementary Figures for "Soils and sediments host novel archaea with divergent monooxygenases implicated in ammonia oxidation": Supplementary_Fig6_v2.pdf

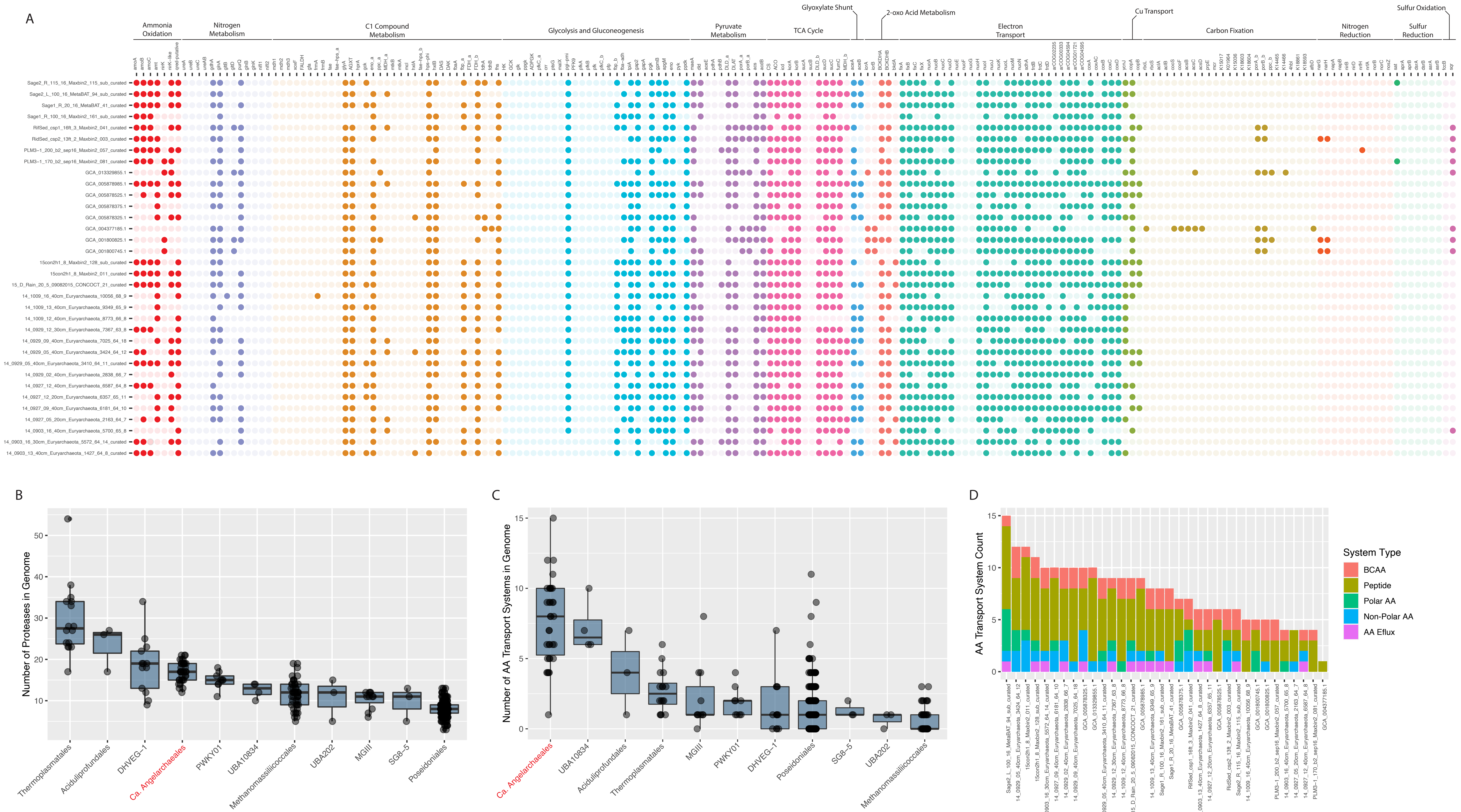


gene names are sometimes repeated if they contribute to mutliple metabolic groups or functional guilds and have _a or _b appended to their names
