## Supplementary figures and images for "Soils and sediments host novel archaea with divergent monooxygenases implicated in ammonia oxidation"

### Supplementary_Fig1_v1.pdf

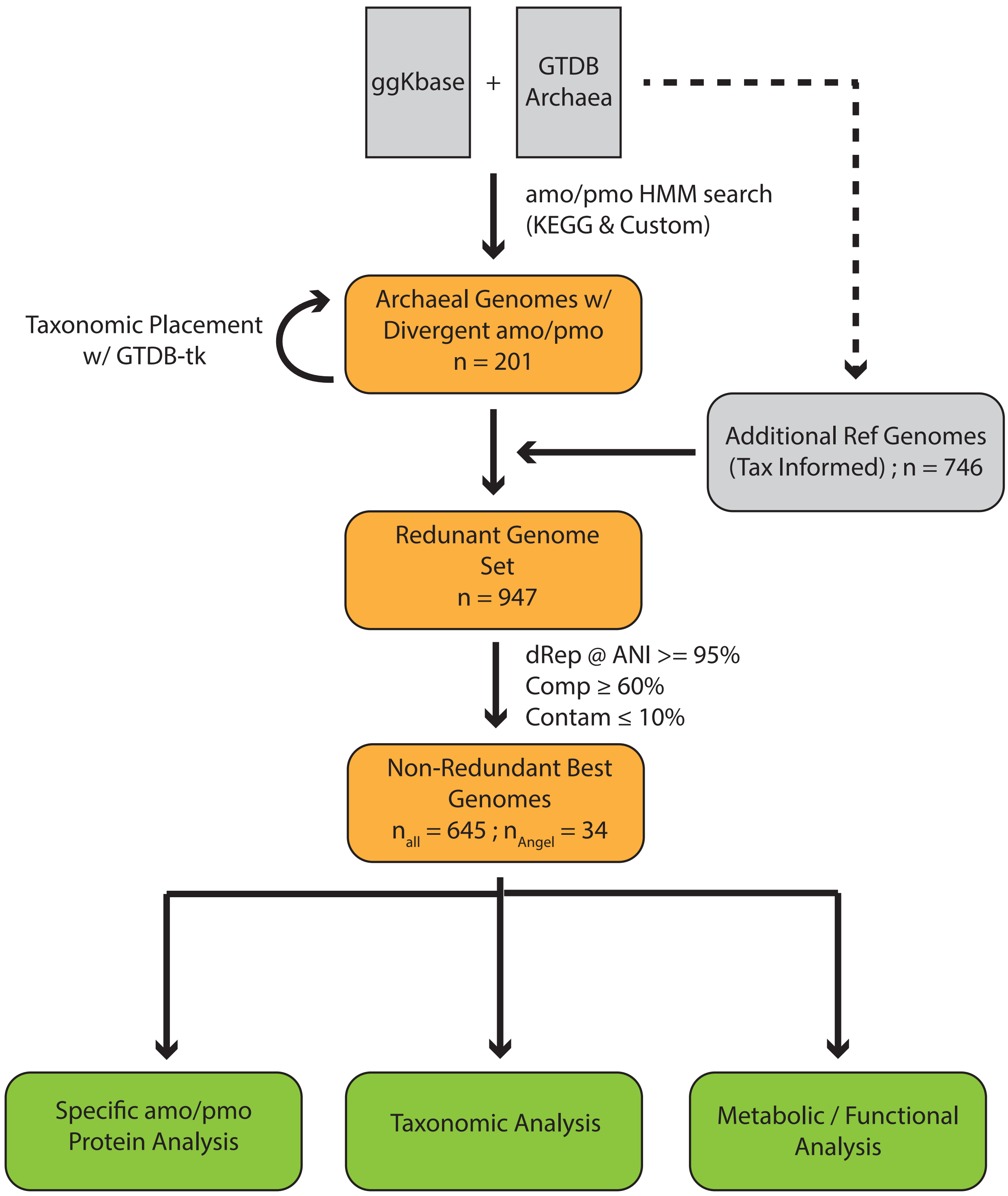

### Supplementary_Fig2_v1.pdf

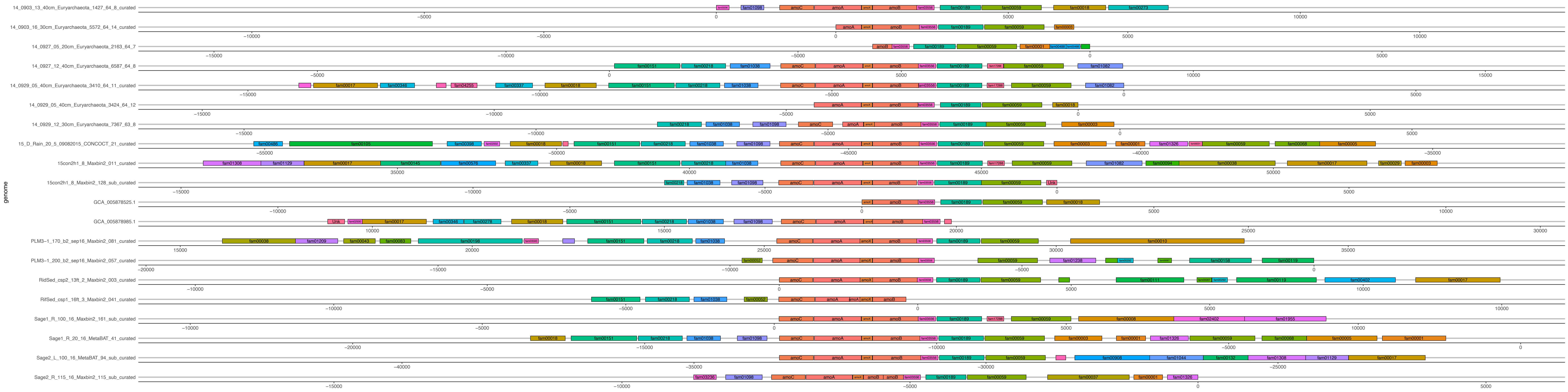

### Supplementary_Fig3_v1.pdf

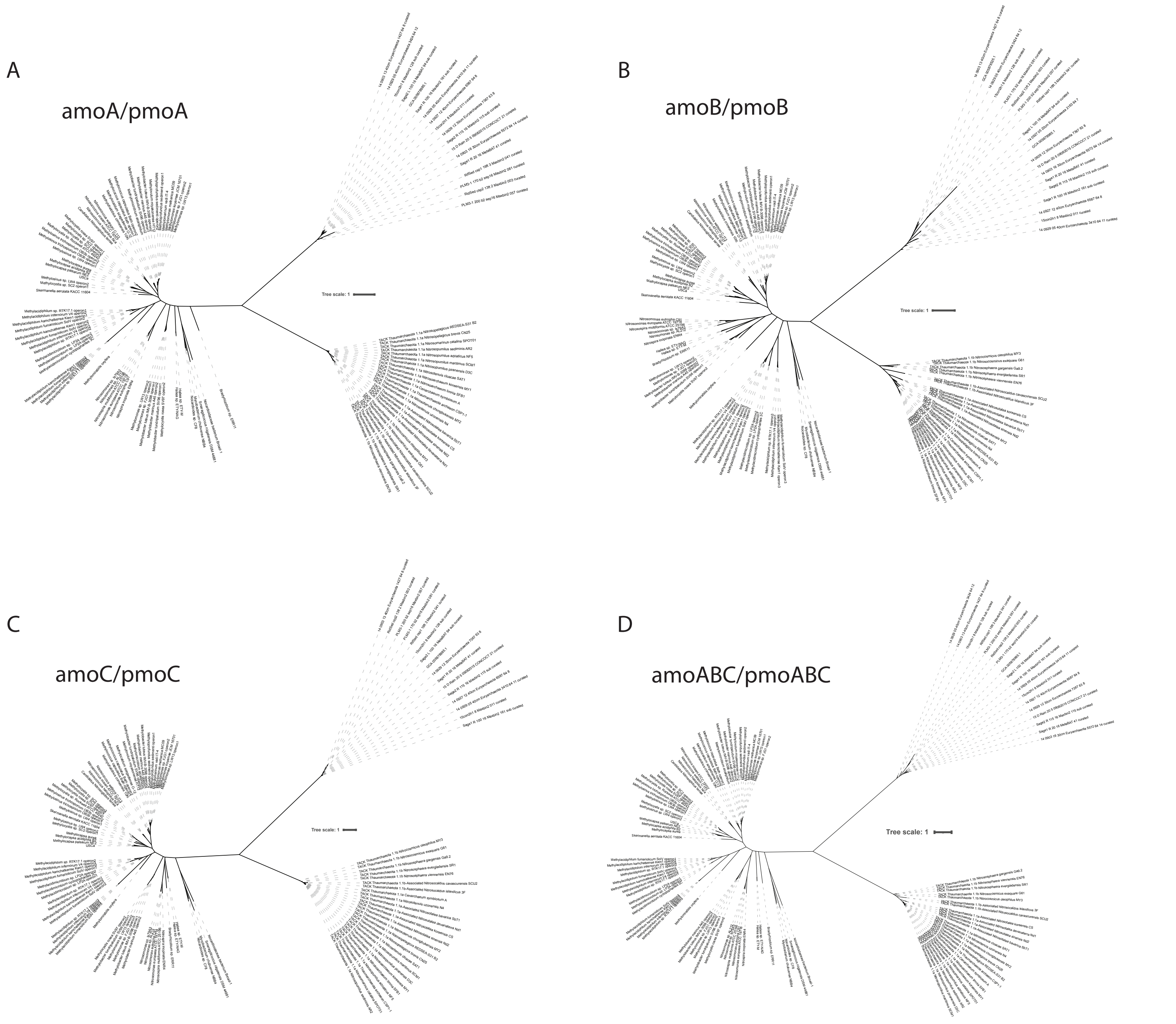

### Supplementary_Fig4_v2.pdf

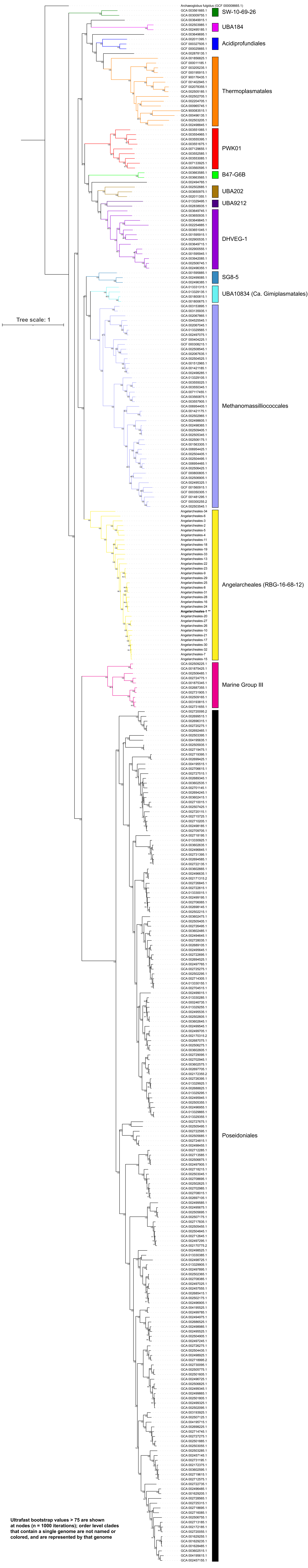

### Supplementary_Fig5_v1.pdf

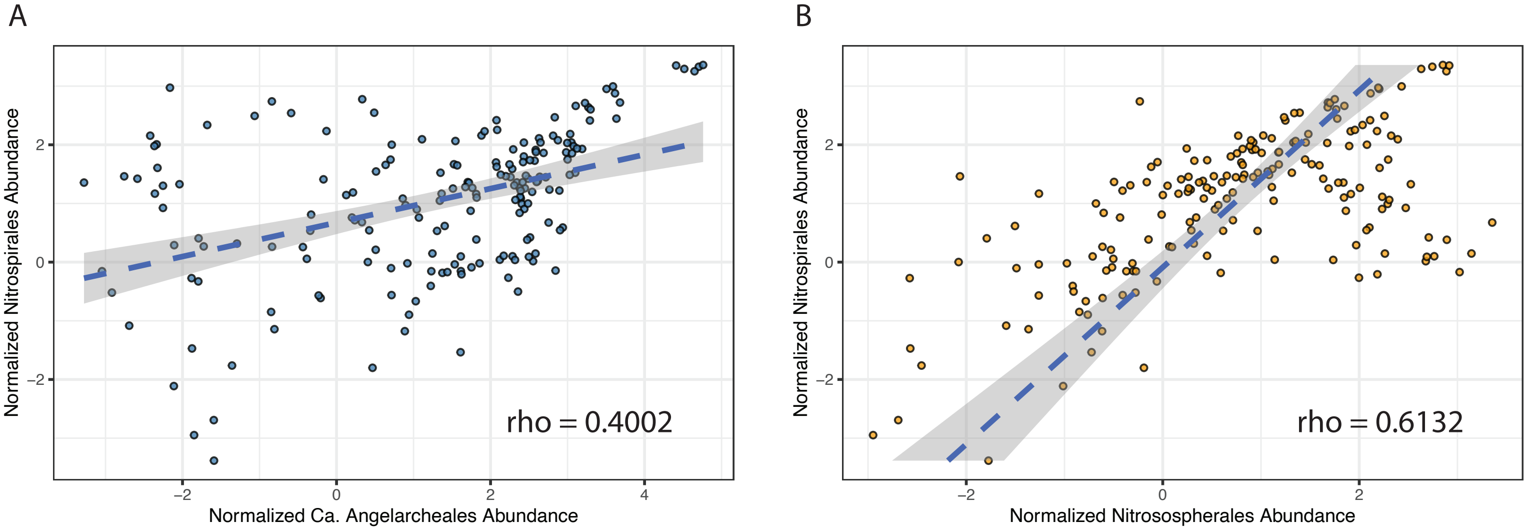

### Supplementary_Fig7_v1.pdf

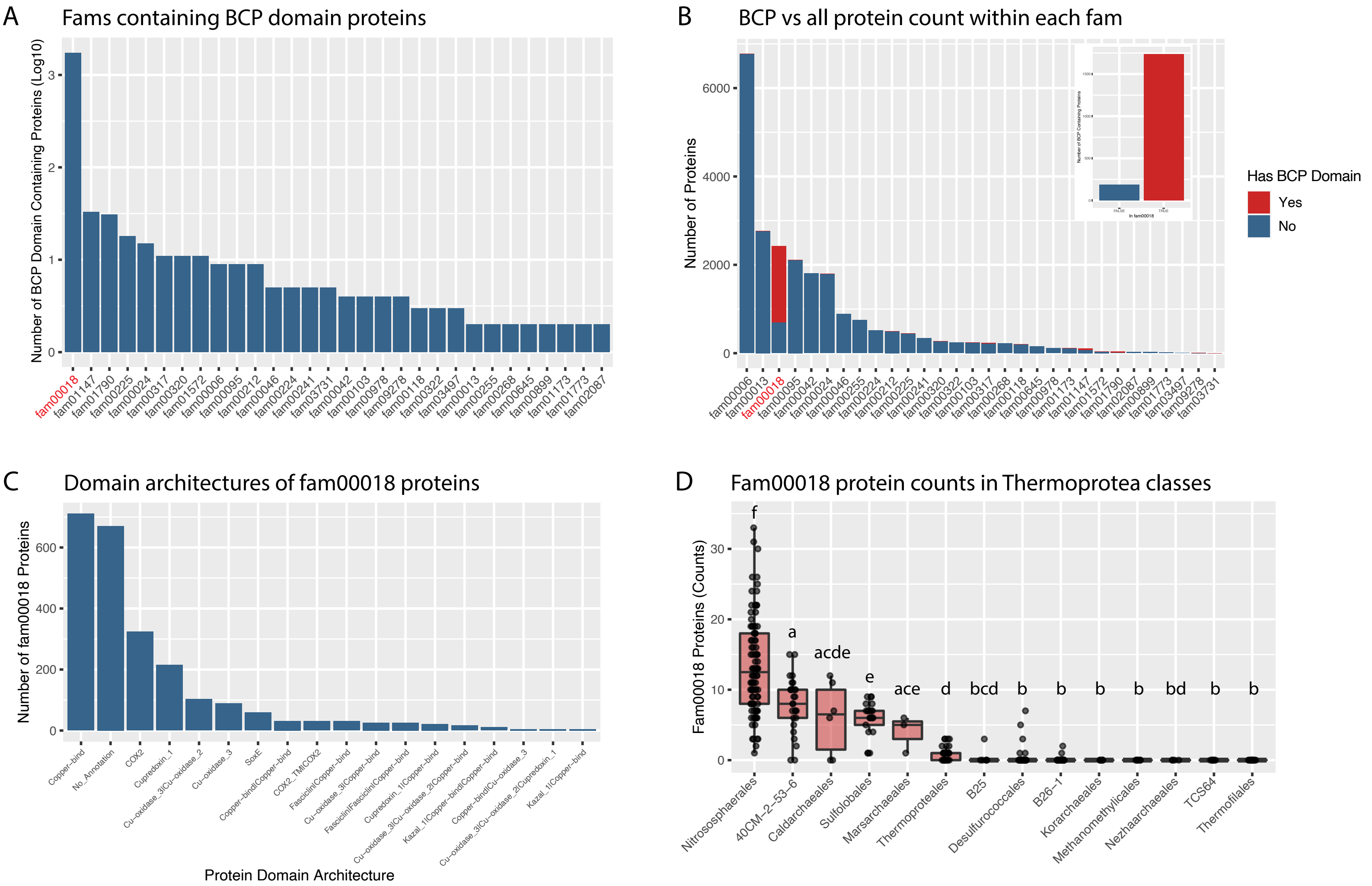

### Supplementary_Fig8_v1.pdf

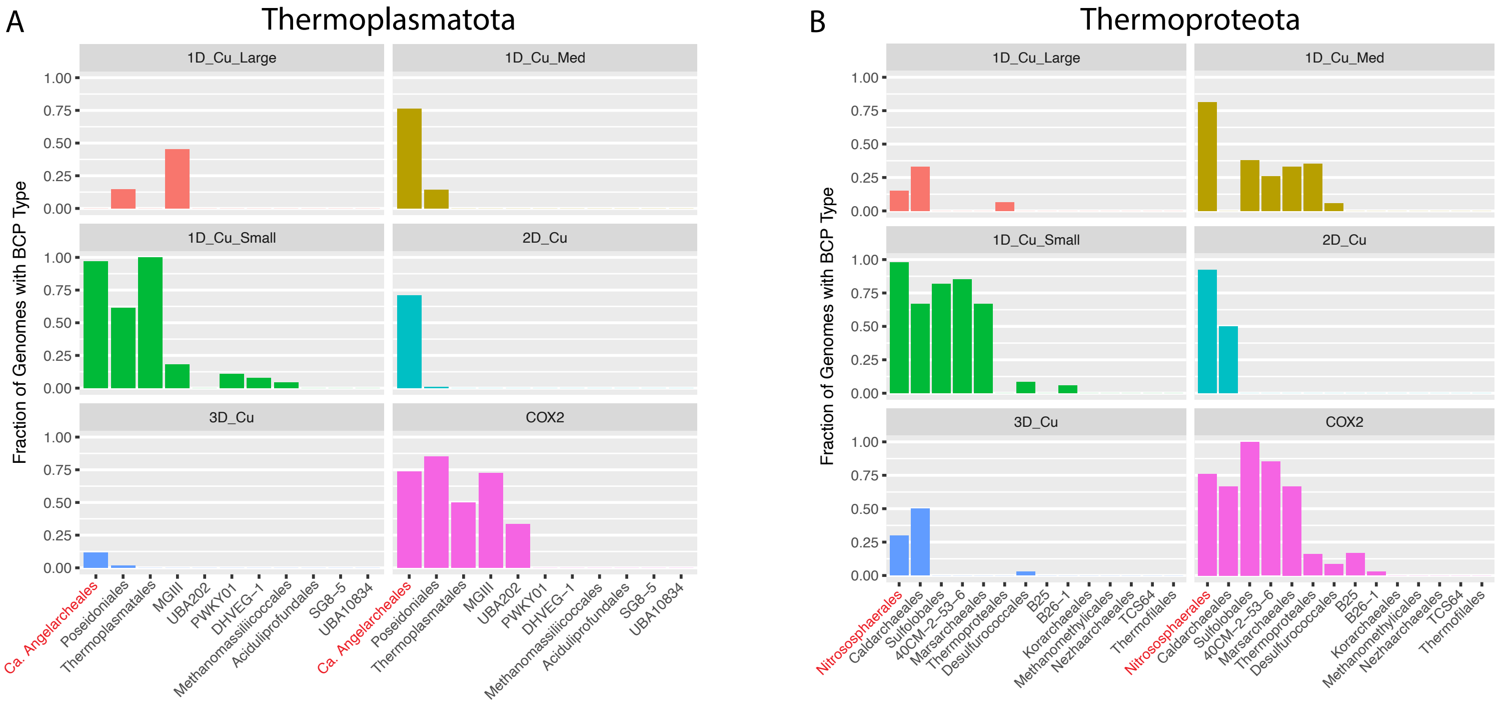

### Supplementary_Fig9_v1.pdf

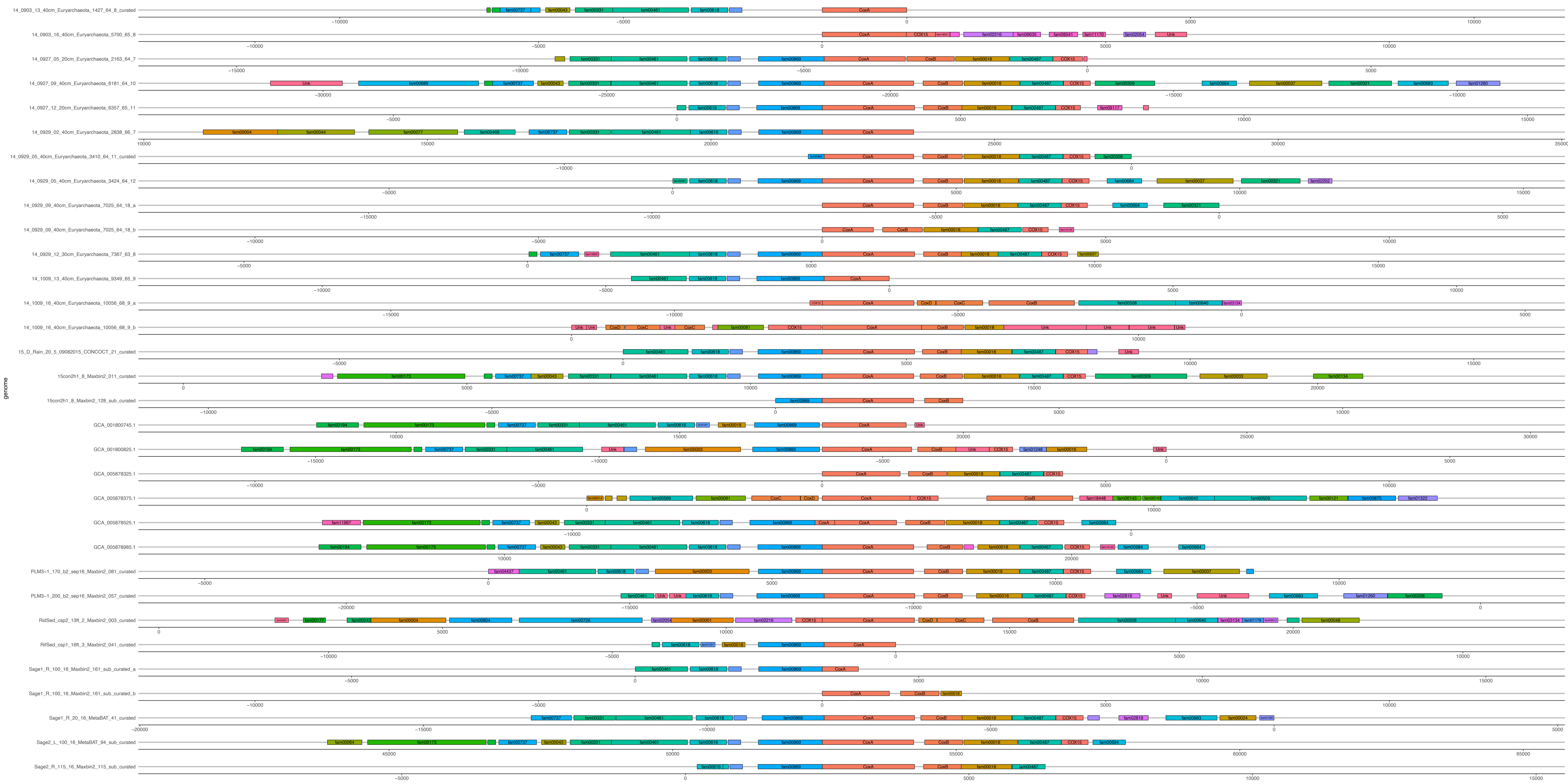

### Supplementary_Fig10_v1.pdf

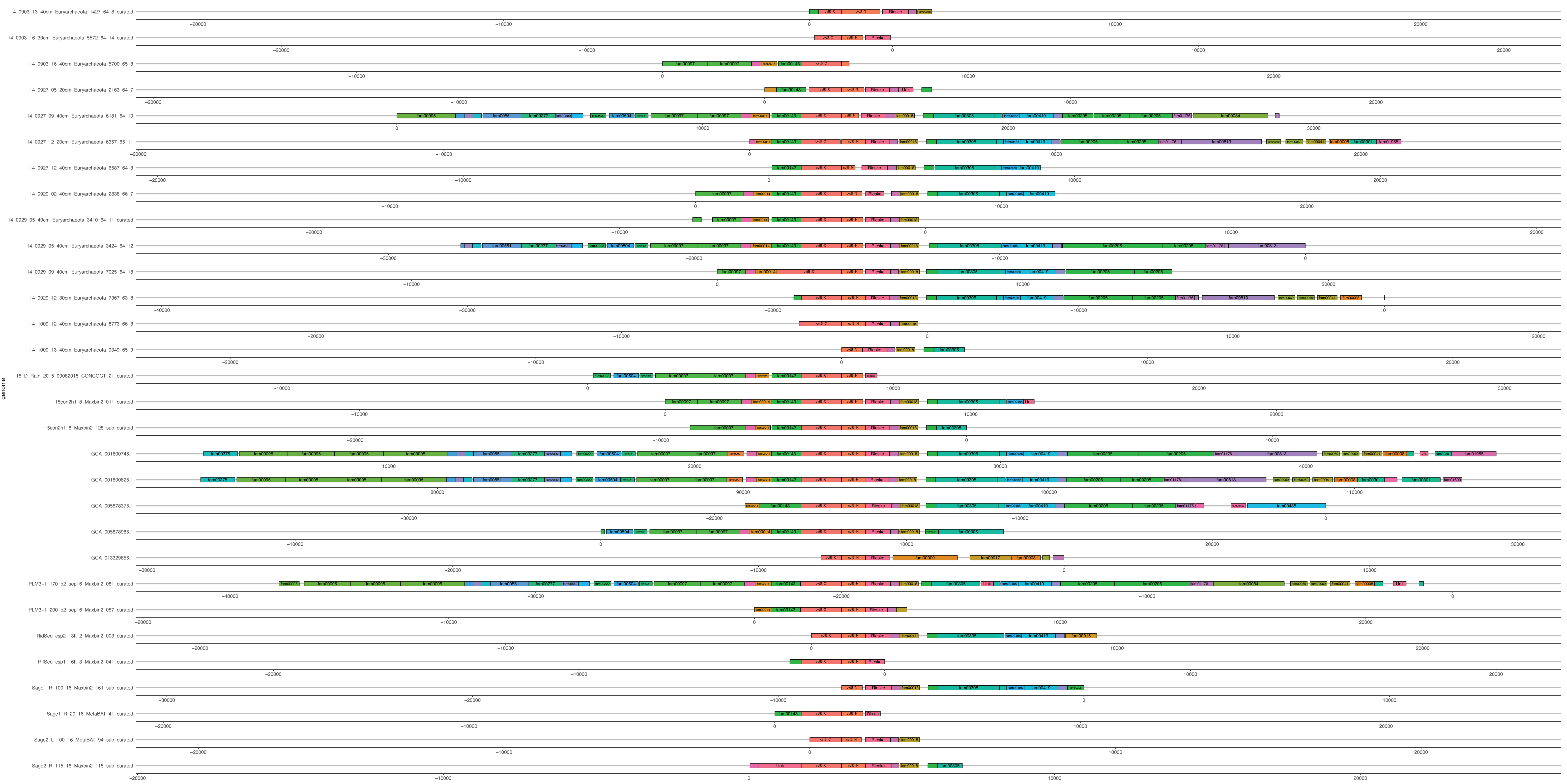

### Supplementary_Fig11_v1.pdf

#### TMHMM Analysis for Proteins of Subfam17112

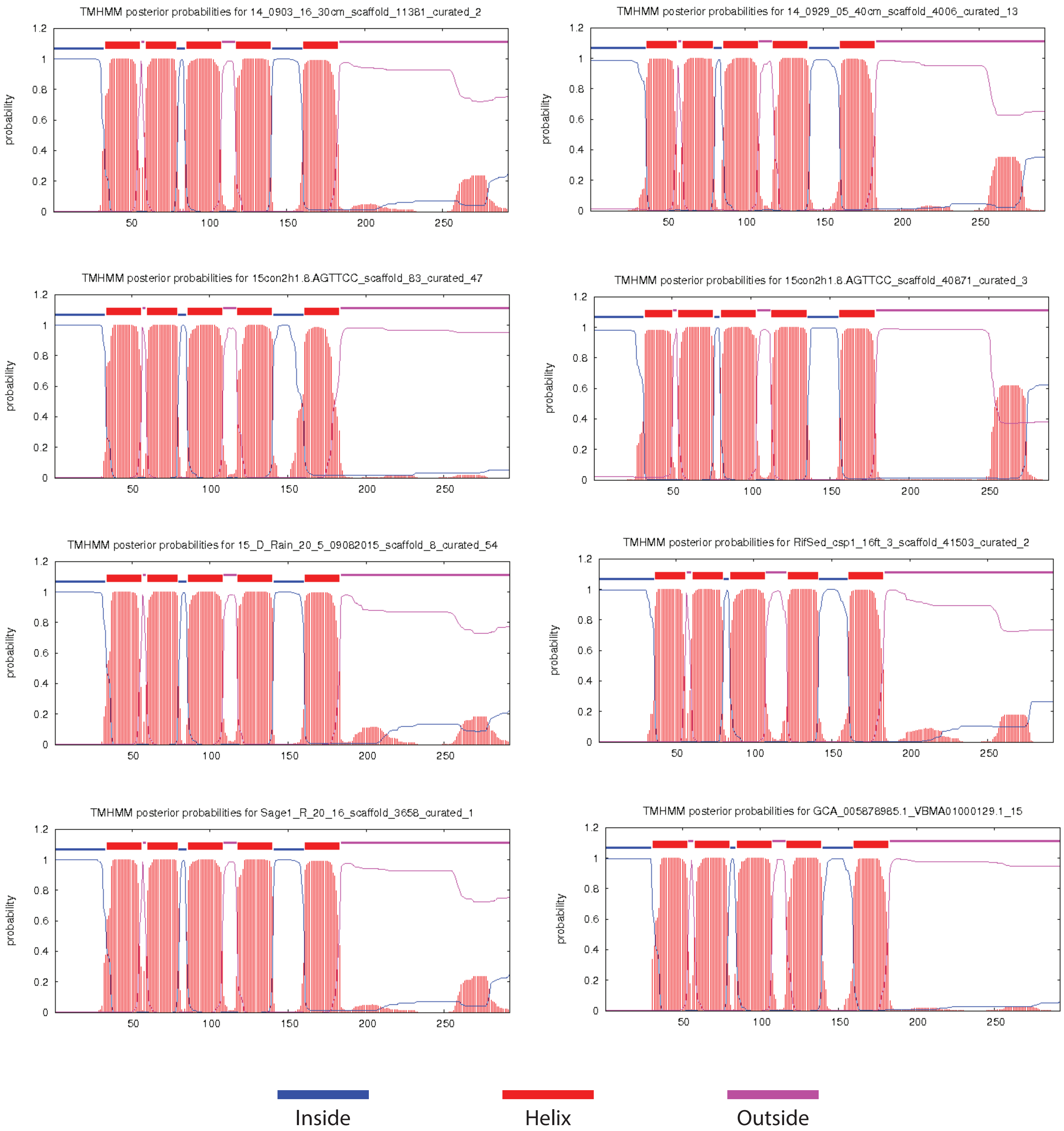
