## Supplementary Figure Legends for "Soils and sediments host novel archaea with divergent monooxygenases implicated in ammonia oxidation"

**Supplementary Figure 1 | Diagram of informatic search strategy.** This flow diagram illustrates the path from initial novel amoA/pmoA sequences to the recovery of the full genome set used for the analysis in this manuscript. Grey boxes indicate databases. Orange boxes indicate sets of genomes with the number of genomes in the set being indicated within the box. Green boxes indicate downstream analysis steps. Black arrows indicate an applied informatics method (noted in figure) or merge of genome sets.

**Supplementary Figure 2 | Diagram of amoCAXB containing contigs.** This figure displays all amoCAXB containing contigs identified in Ca. Angelarcheales in this study. Contigs are centered on the amoB/pmoB gene. All contigs are at the same scale and scales are in base pairs, but scales are arbitrary due to centering of the contigs. The amoCAXB genes in each gene are colored red. Other genes are labeled by their protein family cluster membership, and genes in the same family share the same color. Color for each family is arbitrarily chosen. Genome names where each contig originates are noted on the left of the figure.

**Supplementary Figure 3 | All amo/pmo gene trees. (A)** The maximum likelihood tree for all amoA/pmoA protein sequences from Ca. Angelarcheales and known references (n = 112 sequences). **(B)** The maximum likelihood tree for all amoB/pmoB protein sequences from Ca. Angelarcheales and known references (n = 114 sequences). The maximum likelihood tree for all amoC/pmoC protein sequences from Ca. Angelarcheales and known references (n = 110 sequences). **(D)** The maximum likelihood tree for all concatenated amoABC/pmoABC protein sequences from Ca. Angelarcheales and known references (n = 112 sequences; > 2 subunits per genome) for comparison. All trees are displayed unrooted. Branch names indicate the genome encoding the sequence, and the operon number the sequence originates from if multiple amo/pmo sequences are found in a genome. Tree scales indicate the number of average substitutions per site.

**Supplementary Figure 4 | Full concatenated marker gene phylogenetic tree for Thermoplasmatota phylum.** Maximum likelihood phylogenetic tree constructed for the archaeal phylum Thermoplasmatota using a concatenated alignment of 76 archaeal specific marker genes. The tree includes 32 Ca. Angelarcheales genomes and 295 reference genomes. Colored bars and matching colored branches indicate order level lineages within the tree which are named, with colors being arbitrarily chosen. Order level lineages that contain a single genome were not named or colored. The type strain, Ca. Angelarcheales-1, is marked with two asterisks. Tree was rooted using *A. fulgidus* (GCF_000008665.1) as an outgroup. Bootstrap values >= 75% are shown at branches and calculated using ultrafast bootstrapping (ufboot; n = 1000).

**Supplementary Figure 5 | Relative abundance relationships between orders of archaea and Nitrospirales. (A)** Centered log-ratio normalized abundance data, based on rpL6 read counts, for Ca. Angelarcheales (x-axis) and Nitrospirales (y-axis) across 185 shotgun metagenome samples from 6 sites. **(B)** Centered log-ratio normalized abundance data, based on rpL6 read counts, for Nitrososphaerales (x-axis) and Nitrospirales (y-axis) across 185 shotgun metagenome samples from 6 sites. Each dot represents a single sample site of the 185 samples. Best fit lines are plotted using linear regression, and shaded area indicate standard error of the regressions. Rho of values of proportionality (association) between each pair of organisms is displayed on bottom right of plot and significance of both associations was FDR $< 0.001$ using 9999 permutations (permutation test).

**Supplementary Figure 6 | Metabolic features of Ca. Angelarcheales genomes. (A)** Full set of genes used for predicted metabolism pathways for the 32 Ca. Angelarcheales genomes (also see Fig. 2). Filled dots indicate the presence of a gene that executes a specific metabolic function or reaction. Dots are colored based on shared presence in pathways or specific metabolic reactions as described above the figure, and colors are chosen arbitrarily. Gene names and displayed data are duplicated in the figure if a gene is a component of multiple pathways. Gene names are appended with letters in the case of duplication for ease of reading. For complete explanation of metabolic functions and search criteria see Supplementary Table 8 **(B)** Counts of proteases in the genomes of Thermoplasmatota orders containing ≥ 3 genomes. Each dot represents a genome. Boxes represent the first and third quartile of counts with the centerline being the median counts for a group. Whiskers bound 1.5 x interquartile range of counts. Ca. Angelarcheales is labeled red for ease of viewing. **(C)** Counts of 26 amino acid and oligopeptide transport systems in the genomes of Thermoplasmatota orders containing ≥ 3 genomes. See Supplementary Table 14 for transporter targets. Each dot represents a genome. Boxes represent the first and third quartile of counts with the centerline being the median counts for a group. Whiskers bound 1.5 x interquartile range of counts. Ca. Angelarcheales is labeled red for ease of viewing. **(D)** Breakdown of amino acid and oligopeptide transport system types in each Ca. Angelarcheales genome. Bar height represents total counts per genome and bar colors represent the fraction of the total transporters of a specific class. Only the 5 transporter classes that were detected in Ca. Angelarcheales are shown.

**Supplementary Figure 7 | Additional statistics on fam00018 proteins. (A)** Total counts of proteins containing pfam identified blue copper protein (BCP) domains across the 30 protein families where at least 1 protein contained a BCP domain. Counts are shown on a log10 scale so counts in all families can be easily viewed together. Fam00018 is colored red for ease of viewing. **(B)** The total number of proteins clustered into each of the 30 protein families where at least 1 protein contained a BCP. Bar color indicates the fraction of the counts that correspond to proteins with an identifiable BCP domain. Inset; Plot shows the number of proteins that have BCP domains and are in fam00018 (red) and not in fam00018 (blue). **(C)** Counts of proteins within fam00018 that have the domain architecture indicated on the x-axis. Names given are pfam domain names, and if multiple domains are present in a protein they are separated by "|" in the order that they occur. Only domain architectures that occurred in more than 5 proteins are included. Counts in the group "No_Annotation" had no identifiable pfam domains above default cutoffs. **(D)** Counts of fam00018 proteins in genomes from each order level lineage of the

Thermoproteota containing ≥ 3 genomes. Each dot represents one genome. Boxes indicate the first and third quartile of counts, lines in boxes indicate median values, and whiskers indicate 1.5 x interquartile range in either direction. Letters above boxes indicate statistically significant differences between groups. Groups sharing no letters have statistically significant differences (FDR ≤ 0.05; pairwise Wilcoxon test).

**Supplementary Figure 8 | Quantification of manually BCP protein subtypes in genomes. (A)** Fraction of total genomes in each order level lineage of the Thermoplasmatota (containing ≥ 3 genomes) that contain at least one protein in the manually subclassified BCP group noted in each panel. Bars are colored arbitrarily based on group classification. Ca. Angelarcheales is in red type for ease of viewing. **(B)** Fraction of total genomes in each order level lineage of the Thermoproteota (containing ≥ 3 genomes) that contain at least one protein in the manually subclassified BCP group noted in each panel. Bars are colored arbitrarily based on group classification. Nitrososphaerales is in red type for ease of viewing. BCP subtypes were manually classified for 100 fam00018 subfamilies that contained ≥ 5 proteins. For full data on subfamily level classification see Supplementary Table 9.

**Supplementary Figure 9 | Diagram of coxAB containing contigs.** This figure displays all coxAB (oxygen utilizing terminal oxidase) containing contigs identified in Ca. Angelarcheales in this study. Contigs are centered on the coxA gene. All contigs are at the same scale and scales are in base pairs, but scales are arbitrary due to centering of the contigs. The coxAB genes in each gene are colored red, and fam00018 proteins are labeled beige. Other genes are labeled by their protein family cluster membership, and genes in the same family share the same color. Color for each family is arbitrarily chosen. Genome names where each contig originates are noted on the left of the figure.

**Supplementary Figure 10 | Diagram of Complex III containing contigs.** This figure displays all Complex III (cytochrome bc-like) containing contigs identified in Ca. Angelarcheales in this study. Contigs are centered on the gene containing the N-terminal portion of the split cytochrome b (arCOG01721). All contigs are at the same scale and scales are in base pairs, but scales are arbitrary due to centering of the contigs. All genes are labeled by their protein family cluster membership, and genes in the same family share the same color. Color for each family is arbitrarily chosen. Genome names where each contig originates are noted on the left of the figure.

**Supplementary Figure 11 | Transmembrane region predictions for subfam17112.** This figure displays the TMHMM output for each of the 8 proteins that were clustered into subfam17112. Names of proteins that can be referenced in the full annotation table (on Figshare) are on the top of each panel. Predictions for regions of the protein are presented as lines in the top of each plot with intracellular amino acids blue, transmembrane amino acids in red, and extracellular amino acids in purple. Plots below the prediction model show the posterior probabilities of each prediction, which are visible on the y-axis. The x-axis of each plot shows the amino acid position in the proteins for the predictions.
